# Aromatic amino acids in the finger domain of the FMRFamide-gated Na^+^ channel are involved in the FMRFamide recognition and the activation

**DOI:** 10.1101/2023.02.01.526556

**Authors:** Yasuo Furukawa, Iori Tagashira

**Affiliations:** Laboratory of Neurobiology, Graduate School of Integrated Arts and Sciences; Graduate School of Integrated Sciences of Life, Hiroshima University, Kagamiyama 1-7-1, Higashi-Hiroshima 739-8521, Japan

**Keywords:** **keywords** DEG/ENaC, FaNaC, FMRFamide, mutagenesis, dose-response, docking simulation

## Abstract

FMRFamide-gated Na^+^ channel (FaNaC) is a member of the DEG/ENaC family and activated by a neuropeptide, FMRFamide. Structural information about the FMRFamide-dependent gating is, however, still elusive. Because two phenylalanines of FMRFamide are essential for the activation of FaNaC, we hypothesized that aromatic-aromatic interaction between FaNaC and FMRFamide is critical for FMRFamide recognition and/or the activation gating. Here, we focused on eight conserved aromatic residues in the finger domain of FaNaCs and tested our hypothesis by mutagenic analysis and *in silico* docking simulations The mutation of conserved aromatic residues in the finger domain reduced the FMRFamide potency, the extent of which were dependent on the mutated position, suggesting that the conserved aromatic residues are more or less involved in the FMRFamide-dependent activation. The kinetics of the FMRFamide-gated currents were also modified substantially in some mutants. The docking simulations revealed the aromatic-aromatic interaction between some conserved aromatic residues in FaNaC and FMRFamide. Collectively, our results suggest that the conserved aromatic residues in the finger domain of FaNaC are important determinants of the ligand recognition and/or the activation gating in FaNaC.

## Introduction

The degenerin/epithelial sodium channel (DEG/ENaC) family is a diverse family of ion channels which are essential for Na^+^ absorption, mechanotransduction, acid-sensing and peptidergic neurotransmission [24, 25, 28, 35]. The epithelial sodium channels (ENaC) and the acid-sensing ion channels (ASIC) are well studied archetypes in the DEG/ENaC family. The DEG/ENaC channels are homo- or hetero-trimers [26, 37, 52], and their extracellular domains are divided into five subdomains, that is, thumb, finger, knuckle, palm and *β*-ball [26]. The numerous structural and functional studies on ASIC and ENaC show that the initial gating motion of the DEG/ENaC channels seems to occur in the finger and thumb domains [6, 26, 32, 37, 51, 52]. The initial crystallographic study of ASIC has revealed that the acidic pocket in the interface between the finger and the thumb domains is involved in H^+^ detection and activation gating [26]. By comparing the closed and open structures of cASIC1, it is suggested that the motion for the activation gating of ASIC involves the expansion of the acidic pocket, and that the finger and thumb domains come closer during the activation [52]. ENaC is known to be activated by proteolytic cleavage of the finger domain [29] and a recent cryo-EM study shows that the protease cleaves the structures rocked between the finger and thumb domains [37]. The interactions between the finger and thumb domains therefor seem to be essential for the initial activation motion of the DEG/ENaC family channels.

Another interesting branch of the DEG/ENaC family is peptide-gated cation channels. FMRFamide-gated Na^+^ channel (FaNaC) is a first recognized peptide-gated channel, which is activated by a tetrapeptide, FMRFamide [12, 34, 53]. Until recently, FaNaCs have been cloned and analyzed only in molluscs [20, 27, 34, 38]. A recent comparative phylogenetic survey of FaNaC has, however, confirmed rather broad distribution of FaNaC in invertebrates [14]. Some related peptide-gated channels have also been cloned in limited species. Hydra sodium channels (HyNaC) cloned from *Hydra* are nonselective cation channels which are activated by Hydra-RFamides [16, 17, 21]. The myoinhibitory peptide-gated ion channel (MGIC) in *Platynereis* is activated by myoinhibitory peptides identified in *Platynereis* [43]. Several pharmacological and biophysical features including single channel conductance and kinetics, relative permeability, channel block by divalent cations and amiloride are well known in FaNaC as well as in HyNaC [16, 17, 19, 20, 21, 23, 31, 34, 38, 53]. By contrast, the peptide binding site(s) of these peptide-gated channels are not well elucidated. Needless to say, the knowledge about the agonist binding site(s) of the peptide-gated channels is vital to understand the activation gating of the channels, which may includes the interaction of the finger and thumb domains as in other DEG/ENaC channels.

Cottrell’s group has discovered a short stretch of amino acid sequence (RRMYFNN in *Helix aspersa* FaNaC) which is important for FMRFamide sensitivity in FaNaC [11, 13]. In a homology model of FaNaC, the RRMYFNN sequence makes a part of *α*1-helix inthe finger domain, and Niu et al [36] have shown recently that Y131 in the sequence is important for FMRFamide recognition in *Helix aspersa* FaNaC. Y131 is also conserved in *Aplysia kurodai* FaNaC which shows similar FMRFamide sensitivity, but not in *Helisoma trivolvis* FaNaC which is much less sensitive to FMRFamide. Thus, a tyrosine residue in *α*1-helix of FaNaC seems to be one of the site which may be involved in FMRFamide binding. Another interesting information obtained from earlier pharmacological studies using the analogue peptides is that 1st and 4th phenylalanine residues of FMRFamide is essential for its bioactivity in FaNaC [10].

With these pieces of information in mind, we have hypothesized that the aromatic-aromatic interaction between FMRFamide and FaNaC is involved in the activation of FaNaC. In the present study, we tried to test our hypothesis by mutagenic analysis and *insilico* docking simulations focused on the eight conserved aromatic residues in the finger domain of FaNaC. We found that the substitution of the aromatic amino acids by valine changed EC50 of the FMRFamide dose-response relationship of FaNaC as well as the macroscopic kinetics of FMRFamide-gated currents, and that FMRFamide docking profile in the finger domain of the FaNaC model examined by *in silico* docking simulations was also affected by a single mutation of some of the conserved aromatic residues. The present results suggest that some of the conserved aromatic amino acids in the finger domain of FaNaC are key determinants of the FMRFamide dependent activation of FaNaC.

## Materials and methods

### Ethics

All animal experiments were approved by the Hiroshima University Animal Research Committee (No. G20-1), and performed in accordance with the guide lines for the Japanese Association of Laboratory Animal Science and the Animal Experimentation of Hiroshima University.

### Channels

*Aplysia kurodai* FMRFamide-gated Na^+^ channel (AkFaNaC, DDBJ/GenBank/EMBL accession number AB206707) was used in the present study [20]. Details of the plasmid containing AkFaNaC (pSD64TR-AkFaNaC) was described previously [30]. Following mutants were made by QuickChange (Agilent Technologies, Santa Clara, CA, USA) or PrimeSTAR Mutagenesis Basal kit (Takara Bio Inc., Shiga, Japan): Y156V, W167V, W167F, W167Y, F170V, F174V, F176V, F181V, F188V, F188Y, Y189V, Y189F, Y189S, F188VY189V. The coding sequences of mutants were confirmed by sequencing.

### Oocyte preparation and cRNA injection

The wild-type AkFaNaC (WT) as well as mutant channels were expressed in Xenopus *laevis* oocytes and the oocytes were cultured as described previously [19, 30, 31]. Briefly, frogs were anesthetized in 0.15% MS-222 (Sigma-Aldrich, St. Louis, MO, USA). A small incision was made on the abdomen of anesthetized frog and a part of ovary was dissected out. The incision was then sutured and the frog was maintained in a recovery tank. After the recovery, the frog was returned to a home tank. The ovary was digested by 2% collagenase (Wako Chemicals, Osaka, Japan) dissolved in OR2 medium (in mM: NaCl 82.5, KCl 2, MgCl_2_ 1, HEPES 10, pH 7.5) for 1–2 hours. Dissociated oocytes were collected and stage V–VI oocytes were selected. Selected oocytes were injected with 50 nl of cRNA containing solution and incubated at 18 *^◦^*C in ND96 (in mM: NaCl 96, KCl 2, CaCl_2_ 1.8, MgCl_2_ 1, HEPES 10, pH 7.5). cRNA was synthesized by SP6 RNA polymerase using mMESSAGE mMACHINE SP6 kit (Thermo Fisher Scientific, Waltham, MA, USA). cRNA was routinely injected into oocytes at the marginal zone between the animal hemisphere and vegetal hemisphere. After the incubation for 2–4 days, the oocytes were used for electrophysiological recording.

### Electrophysiological recording

Initially, the expression of FMRFamide-gated currents in oocytes was examined by conventional two electrode voltage clamp by using OC-725C (Warner Instruments LLC., Hamden, CT, USA) as described previously [19, 30, 31]. All the data shown in the main text were, however, obtained by cut-open vaseline gap voltage clamp (COVC) by using CA-1B (Dagan Corporation, Minneapolis, MN, USA) because of its superiority as described below. COVC was carried out as described by others previously [46, 47]. Briefly, the top and middle compartments of the COVC chamber were filled with ND96 and the bottom compartment was filled with K-MES (in mM: KOH 100, EGTA 10, HEPES 10, pH was titrated by methansulfonic acid to 7.4). The oocyte membrane in the bottom compartment was permeabilized by 0.3–0.4% saponin to establish electrical access to the inside of oocyte. The permeabilized oocyte was internally dialyzed with K-MES via a glass pipette (*∼*200 *µ*m in diameter) connected to a syringe pump (flow rate was 2 µl/min). The membrane potential of oocytes was measured by a microelectrode of 0.2–1 MΩ filled with 0.5 M NaCl. Holding potential was –50 mV throughout in the present study. The top compartment which isolates a domed membrane of animal hemisphere of oocyte was continuously perfused with ND96 by gravity (*∼*1 ml/min). The inlet tubings from different reservoirs were connected to a 27G needle which was placed within *∼*1 mm from the domed membrane. FMRFamide (Peptide Institute, Osaka, Japan) was dissolved in ND96 and applied by perfusion for 20 sec. Between the successive application of peptide, oocytes were washed by ND96 more than 5 min (usually 5–7 min) to ensure the recovery from desensitization [19, 30]. All experiments were carried out at room temperature (20–25 *^◦^*C).

The practical liquid exchange rate in the present COVC system was estimated by the change in junction potential at the tip of a blunt microelectrode (Supplementary Fig. 1). We measured the change in junction potential when the perfusion of ND96 was changed to 100 mM KCl and vice versa. We checked five points around the domed membrane of oocyte and found that the change in junction potential depends on the position of the microelectrode in the present experimental condition. The time constant of the rising phase of the change in junction potential was in the range of *∼*250–550 msec if the microelectrode was placed near the surface of domed membrane which faced to the liquid stream. The time constant became longer if the microelectrode was placed behind the dome (*∼*1400 msec). The estimated liquid exchange rate was much faster than the previous estimation in our perfusion system for TEVC (10–90% rise time of the change in junctional potential was more than several seconds) [30].

### Data analysis

Digitized data were analyzed with Clampfit (Ver. 6 or 10, Axon Instruments), Origin (Ver. 8 or 10, Originlab, Northampton, MA, USA), and the open source statistical environment, R [40]. To construct the dose-response relationship of the FMRFamide-gated currents, we usually measured the peak current during the 20 sec exposure to FMRFamide-containing solution. In some mutants, however, the rising phase of the FMRFamide-gated current was such that the current peak was not obtained during 20 sec. In such cases, we measured the current just before the end of FMRFamide perfusion.

The dose-response relationship for FMRFamide was usually approximated by Hill equation of the form,

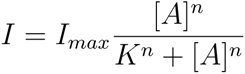

where I is the amplitude of FMRFamide-gated current, I_max_ is the maximum current ofthe dose-response relationship, A is the concentration of FMRFamide, K is EC50, and n is the Hill coefficient.

To compare the steady-state activation of WT and mutant channels, we constructed the normalized dose-response relationships of the channels. The dose-response data obtained in an oocyte was first fitted to the Hill equation, and each data point was then divided by the estimated I_max_. The normalized data were compiled to construct a standard dose-response relationship of the channel. In some mutant channels, the dose-response relationship was better approximated by a sum of two Hill equations.

The macroscopic kinetics of the FMRFamide-gated currents was evaluated by exponential fitting. The sum of two or three exponential function was necessary to approximate the rising as well as the decaying phases of the currents. Because of the liquid exchange rate of our perfusion system described above, the kinetic components with a time constant of less than 1 second were considered to be tentative.

Results of grouped data were expressed as mean*±*SD, and some differences among the channels were examined by multiple comparison (Dunnett’s test). p=0.05 was considered to be a significance level.

### Homology modeling

Structural models of AkFaNaC were made by Modeller ver. 10 [42]. PyMOL [44] and UCSF Chimera [39] were used to make figures. As a template for homology modeling, we used the closed-state cASIC1, 5WKU, [52] obtained from the Protein Data Bank in Europe (http://www.ebi.ac.uk/pdbe/) [49]. The extracellular domain of FaNaC contains FaNaC-specific regions which are absent in ASIC [36]. As shown in our previous modeling [30], the regions are difficult to model appropriately. In the present study, we at first tried to make a model of the full AkFaNaC using 5WKU as a template but the FaNaC-specific regions were not well-defined as before (see Supplementary Fig. 2). Also, the structure of other extracellular subdomains (e.g., the thumb domain) seemed to be affected by the ill-defined FaNaC-specific regions. To make a better model, we removed some of the FaNaC-specific regions (the amino acid number 395–402 and 466–519 of AkFaNaC) which are shown to be dispensable for the FMRFamide-dependent activation of FaNaC [36]. We call this deletion mutant WT-del.

Because a homology between WT-del and cASIC was still low (*∼*25% identity in the sequence used for homology modeling), we placed further restrictions for modeling. Seven conserved S-S bonds, the positions of two *β*-sheets (*β*4, *β*5) and seven *α*-helices (*α*1–7) were pre-determined based on the sequence alignment of ENaC/DEG family channels by Jasti et al [26]. We further placed symmetric restriction on CA atoms as described in (http://salilab.org/archives/modeller usage/2009/msg00084.html). The extracellular structure of WT-del was then well-defined by Modeller. We made 50 models by Modeller and selected a model that showed the least molpdf (the Modeller’s scoring function).

### Docking simulation

To obtain some clues whether the conserved aromatic residues in the finger domain of FaNaC are involved in FMRFamide binding, the ligand docking simulation was done by AutoDock Vina [48]. We used the WT-del model as a receptor (the model is called the WT model in Results and Discussion for simplicity). The structure of FMRFamide cannot be determined explicitly because the peptide is too small. We therefore made the initial structure of FMRFamide by the following method. First, we obtained a putative structure of YGGFMRF using PEP-FOLD3 [33]. We next used Modeller to make a model of FMRF using the YGGFMRF structure as a template. Finally, c-terminal NH_2_ was added to FMRF by hand.

To prepare PDBQT format files of the WT-del and FMRFamide which are used in AutoDock Vina, we used Dock Prep of UCSF Chimera at first. Next, we used a graphic user interface of Chimera to run AutoDock Vina to determine an appropriate docking configuration file for AutoDock Vina. A search area (a rectangle box) was restricted to the upper finger domain of chain A which was large enough to include the conserved aromatic residues (see Fig. 1 and Fig. 9). Size and the coordinate (x,y,z) of the rectangle box were 38*×*30*×*28 Å and (-18.0, 0.0, -50.0), respectively.

**Fig. 1.**
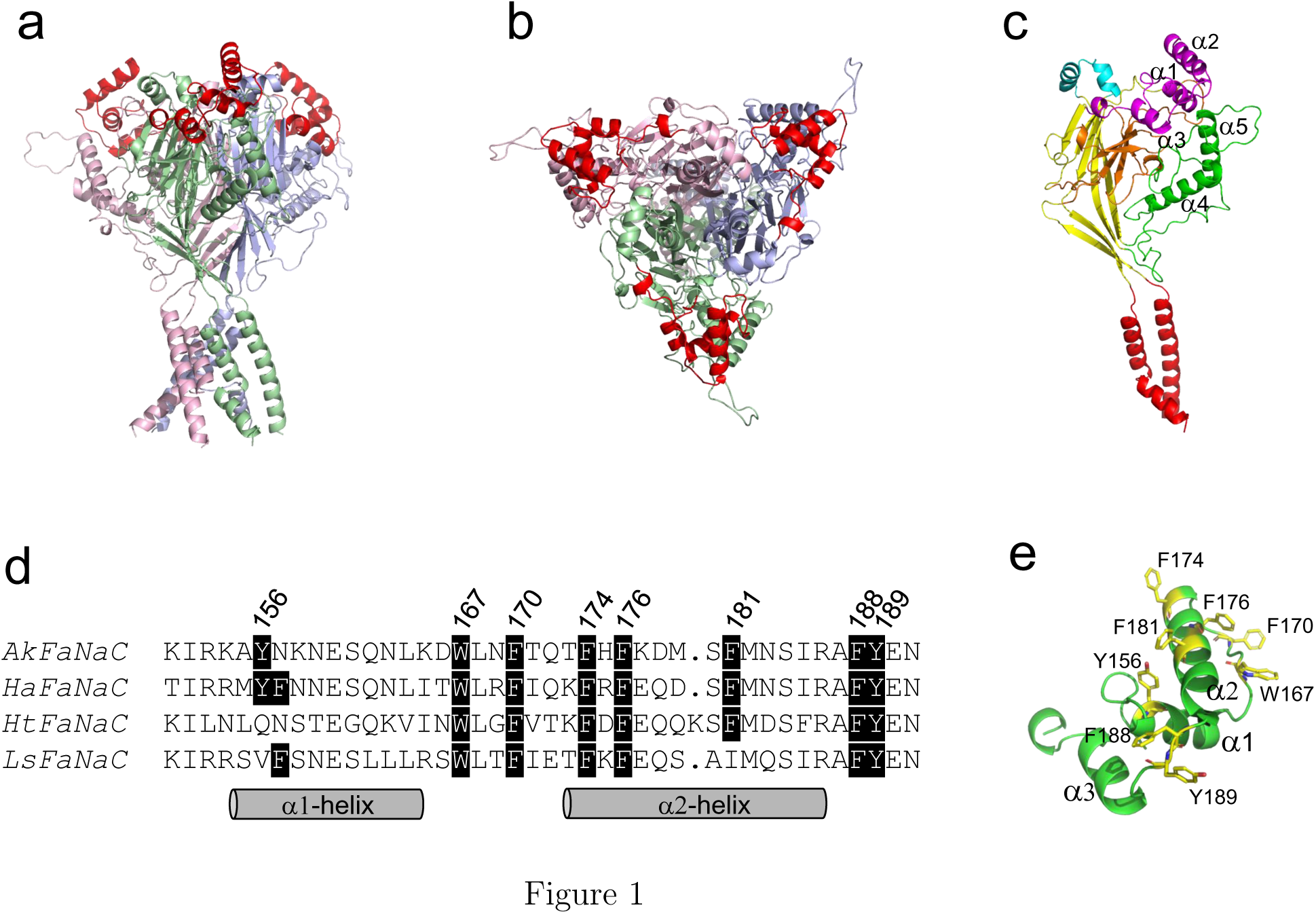
Conserved aromatic residues in the finger domain of FaNaC. **a,b:** A homotrimeric homology model of the deletion mutant of AkFaNaC (WT-del, see Materials and methods for a detail of this model). The finger domain in each subunit is shown by red color. Side view of the model (a). Top view of the model (b). **c:** Subdomains of WT-del. Subdomains are identified and colored according to Jasti et al [26]: thumb (green), finger (purple), knuckle (blue), palm (Yellow), *β*-ball (orange), the transmembrane domains (red). **d:** Sequence comparison of a part of the finger domain among four molluscan FaNaCs. AkFaNaC: *Aplysia kurodai* FaNaC (AB206707), HaFaNaC: *Cornu aspersum* FaNaC (X92113), HtFaNaC: *Planorbella trivolvis* FaNaC (AF254118), LsFaNaC: *Lymnaea stagnalis* FaNaC (AF335548). *Cornu aspersum* and *Planorbella trivolvis* were previously called *Helix aspersa* and *Helisoma trivolvis*, respectively. The numbers shown above are the amino acid number of AkFaNaC. **e:** A part of the finger domain of WT-del showing the conserved aromatic amino acid residues.

By using the docking configuration file as well as the PDBQT files described above, we run AutoDock Vina directly to obtain 2000 docking poses of FMRFamide in the WT-del model. We also carried out similar docking simulations in the mutant channels. The models of mutant channels were made by using a published script in the Modeller Web site (http://salilab.org/modeller/wiki/Mutate%20model) which is an implementation of a method described in Feyfant et al [18].

### Analysis of docking results

A data file made by AutoDock Vina (PDBQT file) contains 3D-coordinates of atoms among others. We first merged all data files (2000 docking results) and extracted 3D-coordinates of *α*-carbons of amino acids in each docking pose. The extraction of necessary information from the merged data file was done by AWK. The resultant text file contains 2000 lines each of which has 3D-coordinates of *α*-carbons of 1st Phe (Phe^1^), Met, Arg and 4th Phe (Phe^4^) of FMRFamide. The reduced data file was then used for further analysis by R [40]. To classify the docking poses efficiently, we used a simple but effective method as described below. At first, 3D-coordinates of a cuboidal space which adequately include the search area of AutoDock Vina was determined. We defined a cuboidal space by a 3D-array having a grid size of 2 Å. In the present case, the 3D-array of 21*×*17*×*15 was prepared for the search area described above. In this case, the total number of unitary cubes was 5355. Because *α*-carbon of each amino acid in FMRFamide is within one of the unitary cubes in the cuboidal space, the position of each *α*-carbon can be specified by a single integer from 1 to 5355, which made the classification of docking poses of FMRFamide much easier. Moreover, because the positions of *α*-carbons of Phe^1^ and Phe^4^ of FMRFamide were found to be sufficient to specify the most poses, the docking poses could be discriminated by just two numbers. The classification methodology was quite efficient to detect similar main chain structures, although there were some fluctuations in the orientations of the main and/or the side chains among the docking poses classified as the same type.

## Results

### Conserved aromatic amino acids in the the finger domain of FaNaC

Figure 1a and 1b show a homology model of a deletion mutant of AkFaNaC and Fig. 1c illustrates a subunit structure of AkFaNaC in which the subdomain structures are shown by different colors as in Jasti et al [26]. The finger domains are situated in the uppermost three corners of the extracellular domain of homo-trimeric structure (Fig. 1a, 1b). The amino acid sequences of a part of the finger domain in four molluscan FaNaCs are shown in Fig. 1d. In the homology model of FaNaC, the sequence identified to be involved in FMRFamide recognition by Cottrell’s group (RRMYFNN in HaFaNaC) is in the *α*1-helix of the finger domain (Fig. 1d). By looking at the nearby sequences of four molluscan FaNaCs, seven other conserved aromatic amino acids are readily noticed (Fig. 1e). Some of these aromatic residues are also well conserved in other putative FaNaCs found in the NCBI database (see Supplementary Fig. 3), many of which have been shown to express FMRFamide-gated currents recently [14]. Because the conserved aromatic residues around the *α*1- and *α*2-helices are prominent in FaNaCs but not in other DEG/ENaC channels and one of them (Y156) has been shown to be involved in the FMRFamide recognition [36], we hypothesized that some or all of the other conserved aromatic amino acids in the finger domain may be involved in the FMRFamide-dependent activation of FaNaC. To test this hypothesis, we made the mutant channels of these aromatic amino acids and examined their responsiveness to FMRFamide.

### FMRFamide-gated currents of saponin-permeabilized oocytes under COVC

FaNaC expresses a slowly activating inward Na^+^ current in response to FMRFamide [20, 27, 34]. In the present study, we employed the cut-open vaseline gap voltage clamp (COVC) to examine the FMRFamide-gated Na^+^ current in *Xenopus oocyte*. COVC is superior to conventional two electrode voltage clamp in *Xenopus oocyte* because it permits faster and more stable voltage clamp. Additional advantage in the present study was an improved perfusion of agonist because the membrane area from which the expressed currents are measured is restricted to a domed membrane (see Materials and methods, Supplementary Fig. 1).

Fig. 2a shows a family of FMRFamide-gated currents of AkFaNaC in response to 1–300 *µ*M FMRFamide obtained by COVC. The initial rising phase as well as the decaying phase of the FMRFamide-gated currents were faster compared to the ones recorded previously by TEVC. This is probably because the practical application speed of FMRFamide is faster in the present experimental condition. The FMRFamide-gated current fully activated within 1–2 seconds by the application of FMRFamide, whereas the current decay after washing out of the peptide took more than a few minutes (see below).

**Fig. 2.**
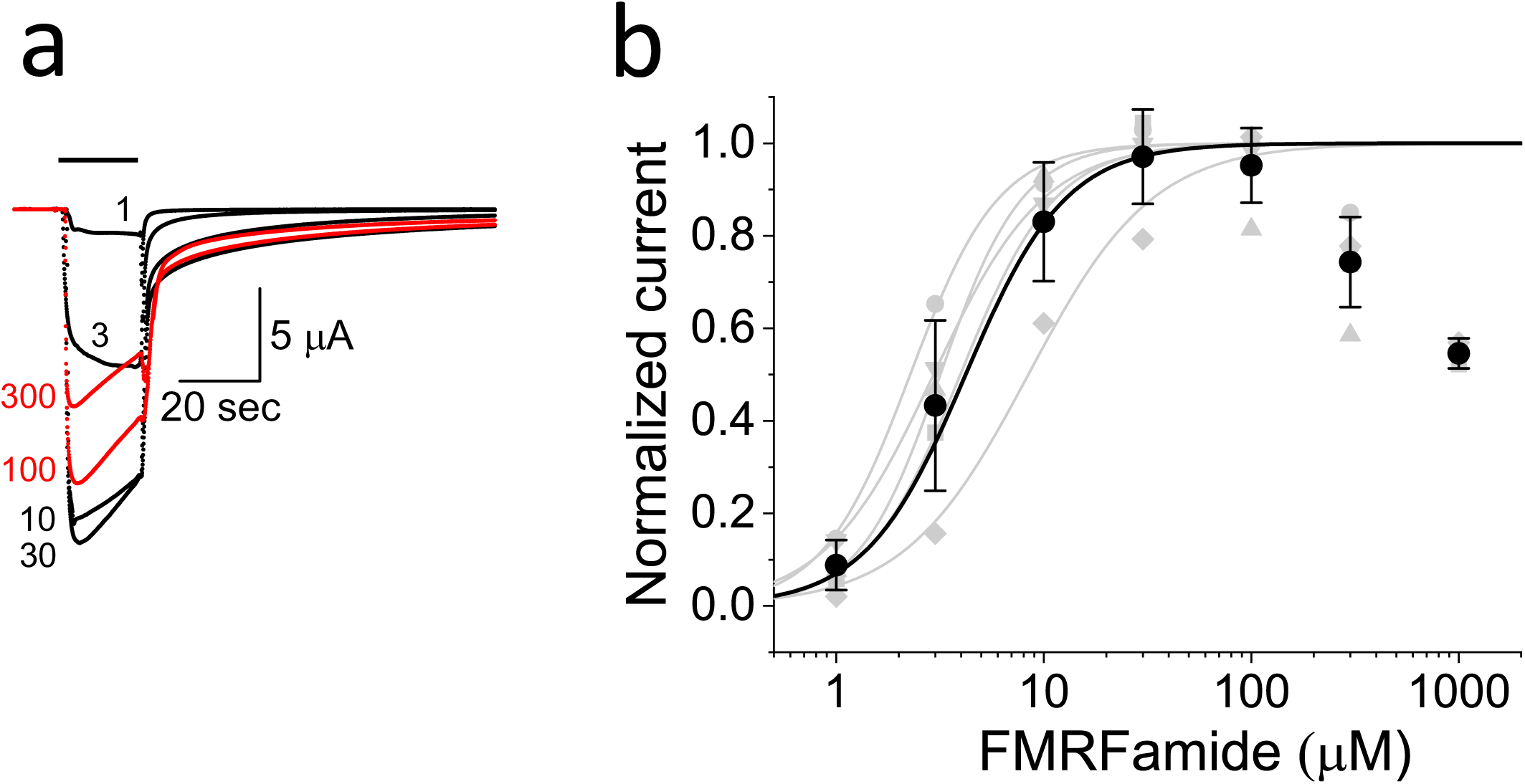
FMRFamide-gated Na^+^ currents in WT. **a:** Example of the family of FMRFamide-gated Na^+^ currents evoked by 20 sec application of FMRFamide indicated by a bar. FMRFamide concentration (in *µ*M) is shown near each trace. The peak currents in WT as well as some mutants show the depression at high concentration of FMRFamide. In this and other figures, such currents are shown by red traces. **b:** The dose-response relationship of WT. Light gray symbols and lines show the dose-response relationships in individual oocytes. Black symbols and error bars are mean*±*SD of the dose-response relationships and a smooth black line is drawn by Hill equation using mean parameters shown in Table 1.

The maximum FMRFamide-gated current of the wild-type AkFaNaC (WT) was usually obtained around the FMRFamide concentration of 30 *µ*M. At higher concentrations, the FMRFamide-gated currents became smaller (see red traces in Fig. 2a), likely due to the desensitization as well as the FMRFamide-induced channel block [22]. When the high concentration of FMRFamide was applied, a transient downward hump was routinely observed in WT as well as some mutants at the beginning of washing out of peptide, suggesting a relief of the agonist-induced pore block [30]. The dose-response relationship up to 100 *µ*M FMRFamide was well approximated by Hill equation (Fig. 2b). Estimated EC50 and Hill coefficient were around 4.2 *µ*M and 1.9, respectively (Table 1). These values were similar to the previously estimated values in our laboratory [19, 20, 30, 31] and those estimated by others [14, 34, 53].

**Table 1.** Dose-response parameters of the wild-type AkFaNaC and the mutants

| channel | EC50 | Hill | EC50.2 | Hill.2 | A1 | A2 | n |
| --- | --- | --- | --- | --- | --- | --- | --- |
| WT | 4.16±2.46 | 1.85±0.30 | – | – | – | – | 5 |
| Y156V | 33.91±17.12<sup>†</sup> | 1.80±0.17 | – | – | – | – | 5 |
| W167V | NA | NA | NA | NA | NA | NA | 5 |
| W167F | 191.31±41.30<sup>‡</sup> | 2.02±0.58 | – | – | – | – | 7 |
| W167Y | 130.45±15.41<sup>‡</sup> | 2.39±0.58 | – | – | – | – | 6 |
| F170V | 54.66±6.66<sup>‡</sup> | 2.23±0.32 | – | – | – | – | 5 |
| F174V | 38.70±5.99<sup>‡</sup> | 1.94±0.44 | – | – | – | – | 8 |
| F176V | 32.25±17.38<sup>†</sup> | 1.46±0.18 | – | – | – | – | 11 |
| F181V | 5.31±1.88 | 1.68±0.39 | – | – | – | – | 6 |
| F188V | 51.04±19.73<sup>‡</sup> | 1.85±0.26 | – | – | – | – | 6 |
| F188Y | 7.66±1.58 | 1.52±0.12 | – | – | – | – | 5 |
| Y189V | 2.50±1.52 | 1.41±0.50 | 475.98±87.37 | 1.43±0.40 | 0.09 ±0.02 | 0.91±0.02 | 5 |
| Y189F | 4.78±0.75 | 1.54±0.30 | 128.68±57.99 | 1.88±0.28 | 0.38±0.04 | 0.62±0.04 | 5 |
| Y189S | 9.04±3.33 | 1.97±0.39 | 144.79±32.57 | 2.60±0.55 | 0.54±0.15 | 0.46±0.15 | 5 |
| F188VY189V | NA | NA | NA | NA | NA | NA | 5 |

The dose-response relationship was fitted by Hill equation as described in Results. Means±SDs of EC50 ( μM) and Hill coefficients are shown. The dose-response relationships of Y189 mutants were approximated by sum of two Hill equations. A1 and A2 are the relative amplitudes of high and low potency components, respectively. n is number of oocytes. W167V and F188VY189V were not activated even by 1 mM FMRFamide.
EC50s and Hill coefficients between WT and the mutants whose dose-response relationships were fitted with a single Hill equation were compared by multiple comparison (Dunnett's test). <sup>†</sup>, $p<0.05$ , <sup>‡</sup>, $p<0.01$ .

### Effects of mutations of the conserved aromatic amino acids in the ***α***1-***α***2 region of the finger domain

We first examined six aromatic residues (Y156, W167, F170, F174, F176, F181) in the α1-helix, α2-helix and a loop between them (see Fig. 1). Aromatic amino acids were substituted to valine and the FMRFamide-dependent activation of the mutant channels was explored. We examined the dose-response relationships of the mutants for FMRFamide as described in WT (Fig. 3, Table 1).

**Fig. 3.**
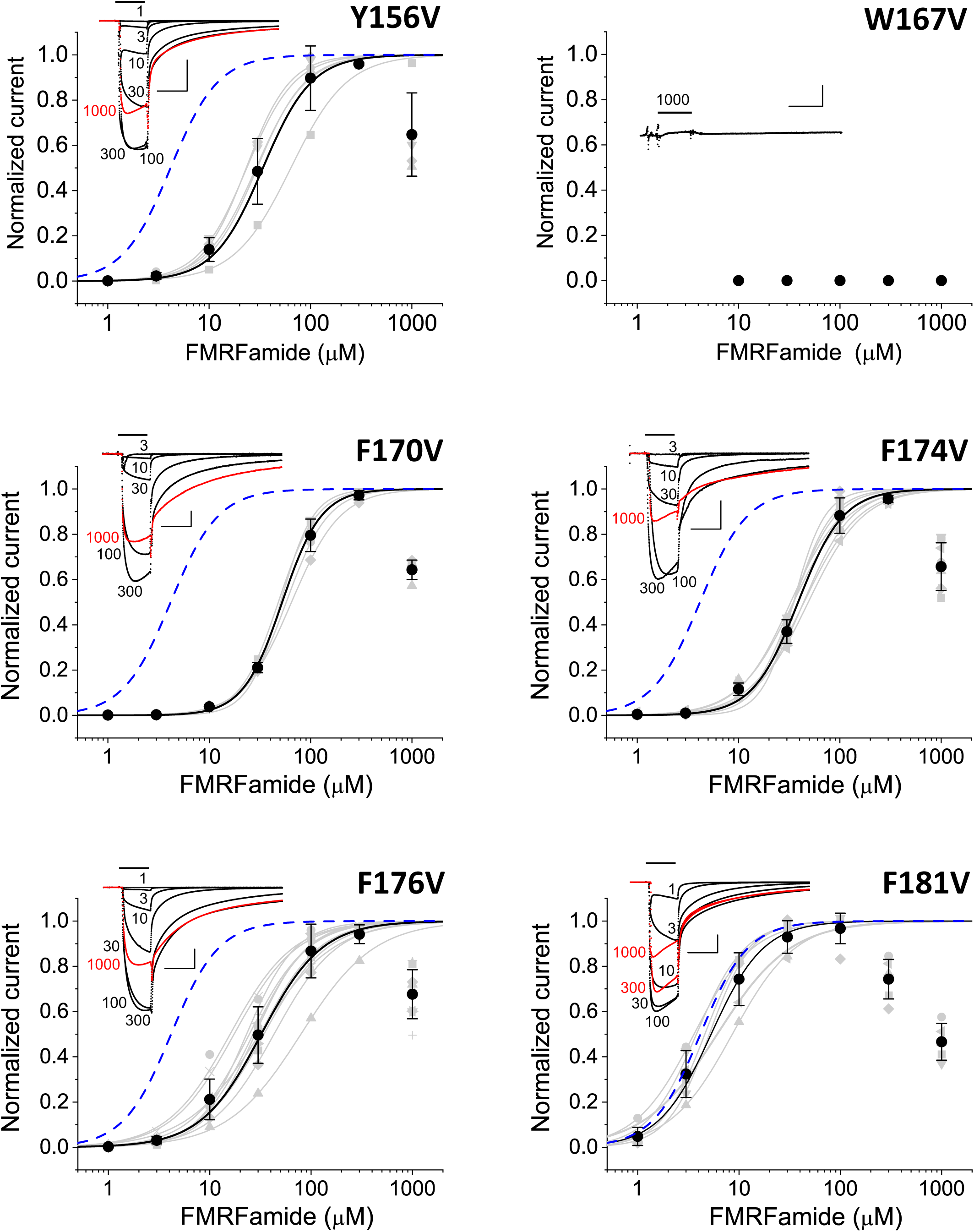
The dose-response relationships of the mutants in the region including *α*1 and *α*2. The inset in each graph shows a family of FMRFamide-gated Na^+^ currents. FMRFamide concentration (in *µ*M) is shown near each trace. Vertical calibrations were as follows: Y156V, 5 *µ*A; W167V, 20 nA; F170V, 300 nA; F174V, 500 nA; F176V, 500 nA; F181V, 2 *µ*A. Horizontal calibrations were 20 sec in all the cases. The dose-response relationships in each graph were made as described in the legend of Fig. 2. The mean EC50s and Hill coefficients for the mutants are shown in Table 1. The blue broken line in each graph indicates the mean dose-response relationship of WT shown in Fig. 2.

Except for W167V, all the tested mutants expressed the FMRFamide-gated currents, indicating that none of these aromatic residues are crucial for FMRFamide-dependent activation. However, EC50s of Y156V, F170V, F174V and F176V but not F181V were *∼*10 fold larger than that of WT, suggesting that these aromatic amino acids are involved in the function of FaNaC. By contrast, FMRFamide did not evoke the current even at 1 mM in W167V, implying that W167 may be critical for the activation of FaNaC.

To address the importance of aromatic moiety at position 167 further, we examined two conservative mutants, W167F and W167Y. As shown in Fig. 4, these mutants were functional, suggesting that the aromatic moiety at position 167 is important for FaNaC.

**Fig. 4.**
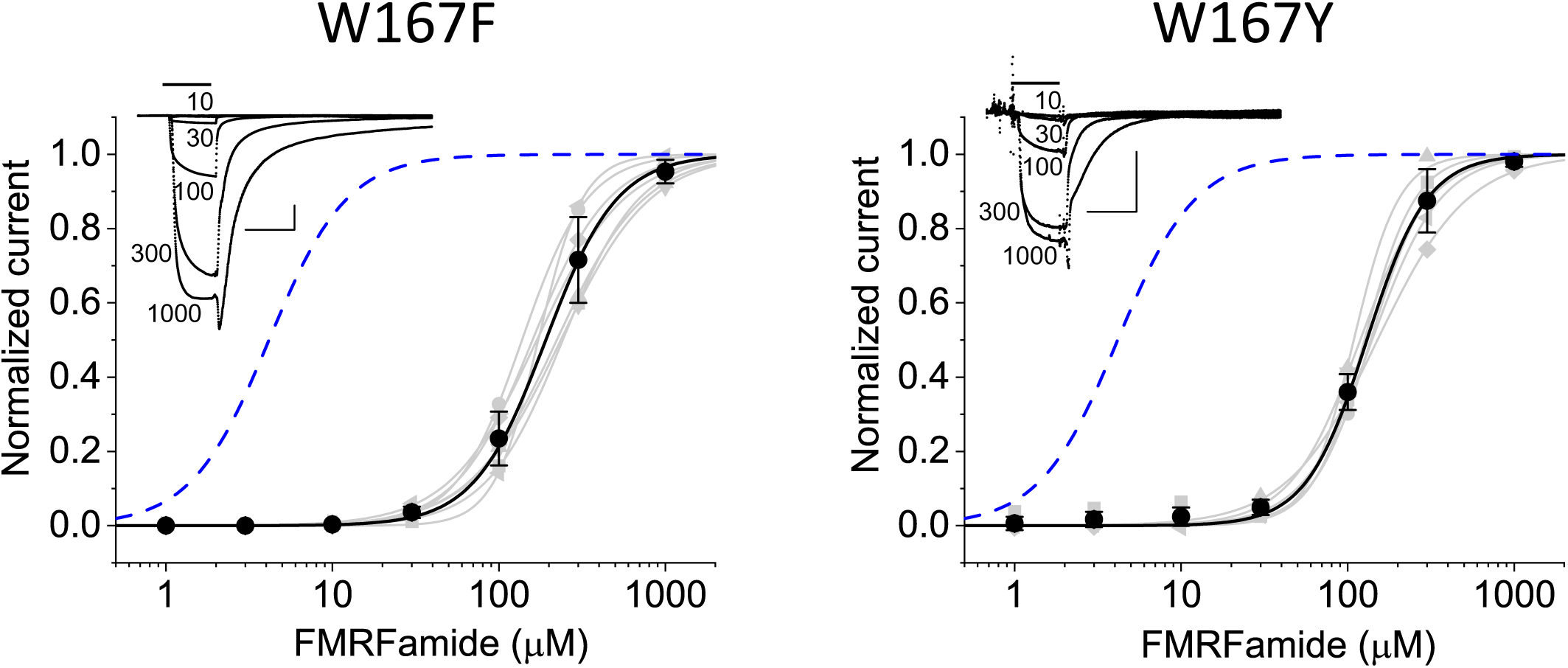
The dose-response relationships of W167F and W167Y. The inset shows a family of FMRFamide-gated Na^+^ currents. FMRFamide concentration (in *µ*M) is shown near each trace. Vertical calibrations for W167F and W167Y were 2 *µ*A and 100 nA, respectively. Horizontal calibration was 20 sec in either case. The dose-response relationships were made as described in the legend of Fig. 2. The mean EC50s and Hill coefficients for the mutants are shown in Table 1. The blue broken lines indicate the mean dose-response relationship of WT.

EC50s of these mutants were, however, much larger than WT and more than 100 µM (Table 1), indicating that the aromatic moiety of phenylalanine or tyrosine at position 167 is not sufficient to reproduce the functionality of FaNaC. These results are consistent with a notion that tryptophan at position 167 is critical for the FMRFamide-dependent activation of FaNaC.

### Effects of mutations of two tandem aromatic amino acids in the loop between *α***2-helix and** *α***3-helix**

We next examined two tandem aromatic amino acids (F188, Y189) in the loop structure between *α*2-helix and *α*3-helix (see Fig. 1). The dose-response relationships of F188 and Y189 mutants for FMRFamide are illustrated in Fig. 5, and EC50s and Hill coefficients are summarized in Table 1. Both F188V and Y189V mutants were activated by FMRFamide but the apparent FMRFamide potency was much less than that of WT. The EC50 of F188V was *∼*10 times larger than that of WT, suggesting that F188 is involved in the activation of FaNaC. Y189V was the least responsive mutant among the tested mutant channels except for non-functional ones. The dose-response relationship of Y189V was not well-fitted with a single Hill equation but approximated by a sum of two Hill equations (Table 1). Interestingly, EC50 for the high sensitive component was rather similar to that of WT. The very low pontency of FMRFamide in Y189V resulted from the fact that the relative amplitude of the low sensitive component was much larger.

**Fig. 5.**
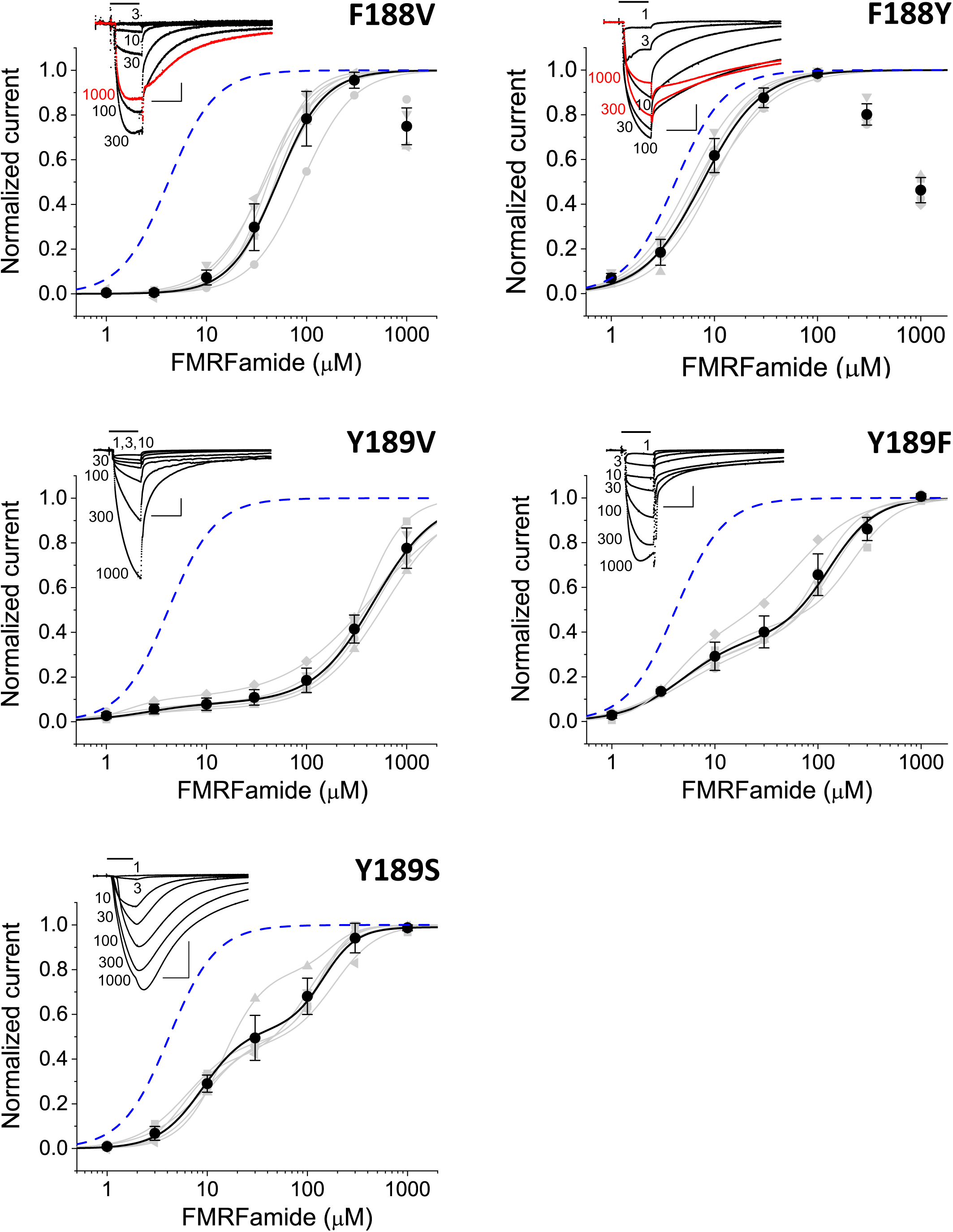
The dose-response relationships of the mutants in the loop between *α*2 and *α*3. The inset in each graph shows a family of FMRFamide-gated Na^+^ currents. FMRFamide concentration (in *µ*M) is shown near each trace. Vertical calibrations were as follows: F188V, 200 nA; F188Y, 200 nA; Y189V, 300 nA; Y189F, 200 nA; Y189S, 500 nA. Horizontal calibrations were 20 sec in all the cases. The dose-response relationships in each graph were made as described in the legend of Fig. 2. The mean EC50s and Hill coefficients for the mutants are shown in Table 1. The blue broken line in each graph indicates the mean dose-response relationship of WT.

The FMRFamide-gated current of Y189V activated quite slowly and did not reach stationary level by 20 sec peptide application especially at high concentrations. Also, the dose-response relationship did not saturate even at 1 mM FMRFamide. Although such features of this mutant hindered the estimation of dose-response parameters, EC50 for the low sensitive component in Y189V should be certainly more than several hundred µM. Because F188V as well as Y189V were functional, neither F188 nor Y189 is indispensable. However, because they are tandem residues, the aromatic moiety of one of them may be good enough to maintain some functionality of FaNaC. To check this possibility, we made a double mutant (F188VY189V) and examined its responsiveness to FMRFamide. F188VY189V was not activated even by 1mM FMRFamide (Table 1), suggesting that the aromatic moiety at position 188 or 189 is necessary for the activation of FaNaC.

To see the importance of aromatic moieties at these sites, we next examined two conservative mutants, F188Y and Y189F. The dose-response relationship of F188Y was close to that of WT (Fig. 5, Table 1), consistent with a notion that an aromatic moiety at position 188 is important for the activation of FaNaC. The macroscopic kinetics of F188Y was, however, quite different from WT (compare the F188Y currents in Fig. 5 and the WT currents in Fig. 1), suggesting that an extra hydroxyl group at position 188 modulates the gating kinetics of FaNaC. Compared to Y189V, the macroscopic activation kinetics of Y189F was close to that of WT (Fig. 5). The dose-response relationship of Y189F was still comprised of two components as in Y189V, but the relative amplitude of the high sensitive component became larger compared to Y189V (Table 1). Apparently, the overall FMRFamide responsiveness of Y189F was better than Y189V, suggesting that the aromatic moiety at this position is also important for the function of FaNaC.

Because a conservative mutant, Y189F, still showed a substantial alteration of the dose-response relationship compared to WT, we speculated that a hydroxyl group of tyrosine might be important. To test this possibility, we replaced Y189 with serine which has an aliphatic hydroxyl group. Both the rising and decaying phases of the FMRFamide-gated currents in Y189S became much slower than that of WT (Fig. 5), again suggesting the importance of aromatic moiety at position 189 for FaNaC. Although the slowly activating currents undermine the analysis of dose-response relationship of Y189S as in Y189V, the dose-response relationship of Y189S clearly showed two components and the overall responsiveness was improved compared to Y189V as in Y189F (Fig. 5, Table 1). The results suggest that a hydroxyl group of Y189 is also important for the function of FaNaC.

### Comparison of the macroscopic kinetics

The FMRFamide-gated current of WT activates within a few seconds but decays much more slowly (see Fig. 2). To quantify the macroscopic kinetics of FaNaC, the rising and decaying phases of the FMRFamide-gated currents were approximated by exponential functions. The fitting results of WT as well as the mutants are illustrated in Figs. 6–8, and fitting parameters obtained from almost fully activated channel currents are compared in Tables 2 and 3. In the following, we describe the rising phase as activation and the decaying phase as deactivation, although both phases are actually the amalgam of transitions among several distinct states of the channels (ex., closed, opened, desensitized, etc).

**Fig. 6.**
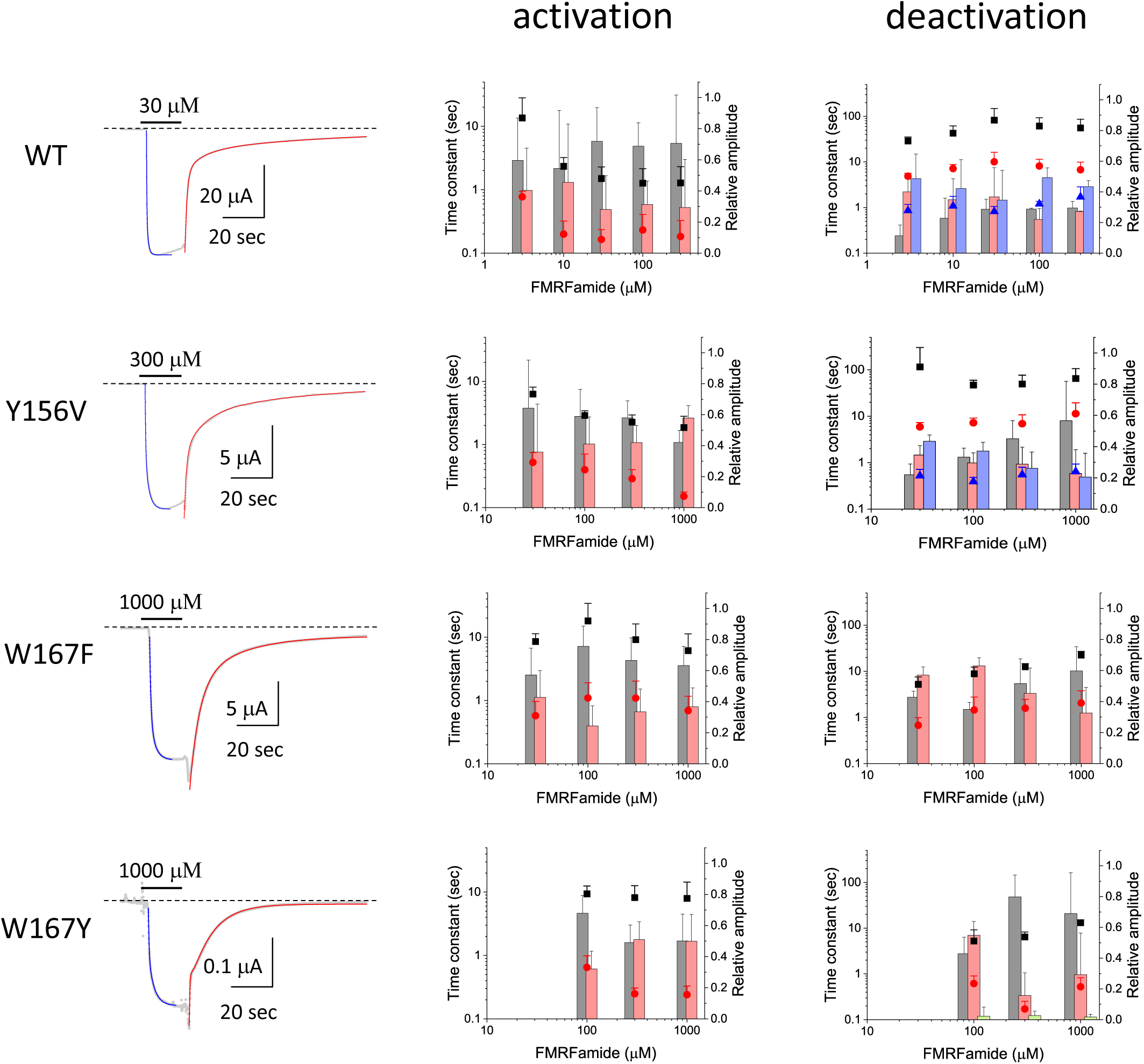
The macroscopic kinetic analysis of WT, Y156V, W167F and W167Y. The rising phase of the FMRFamide-gated Na^+^ current (activation) was approximated by two exponential function with a constant and the decaying phase of the current (deactivation) was approximated by two or three exponential function without a constant except for W167Y. A constant component was necessary to better fit the current decay of W167Y with exponential functions. The left column of this figure shows examples of the fitting. FMRFamide was applied for 20 seconds as indicated by a bar. The broken lines show the zero level. Blue lines indicate the exponential functions fitted for the activation and the red lines indicate the exponential functions fitted for the deactivation. The middle and the right columns show the time constants as well as the relative amplitudes of the fitted exponentials for the activation and the deactivation, respectively. The slow and fast time constants are shown by black squares and red circles, respectively. The relative amplitudes of the slow and fast components are shown by gray and pink bars, respectively. The intermediate time constants and their relative amplitudes, if exists, are shown by blue triangles and light blue bars, respectively. Green bars in the deactivation graph of W167Y show the relative amplitudes of constant components. Number of oocytes was as follows: WT (5), Y156V (5), W167F (7), W167Y (4–6).

**Fig. 7.**
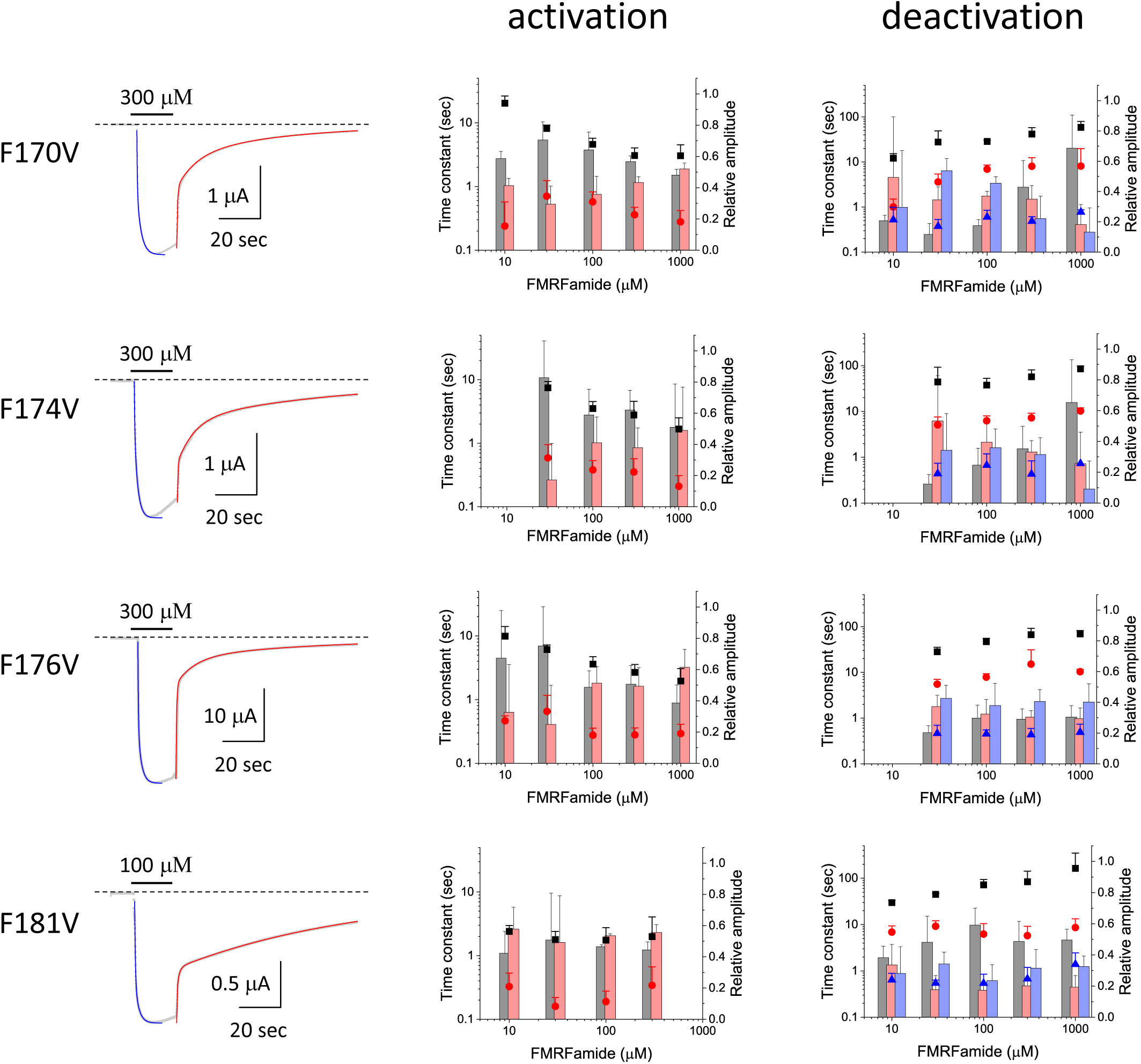
The macroscopic kinetic analysis of F170V, F174V, F176V and F181V. The activation was approximated by two exponential function with a constant and the deactivation was approximated by three exponential functions without a constant. The left column shows examples of the fitting. Superimposed blue and red lines are fitted exponential functions. The meanings of symbols and bars for the activation (middle column) and for the deactivation (right column) are as described in the legend of Fig. 6. Number of oocytes was as follows: F170V (4), F174V (5), F176F (8), F181Y (4).

**Fig. 8.**
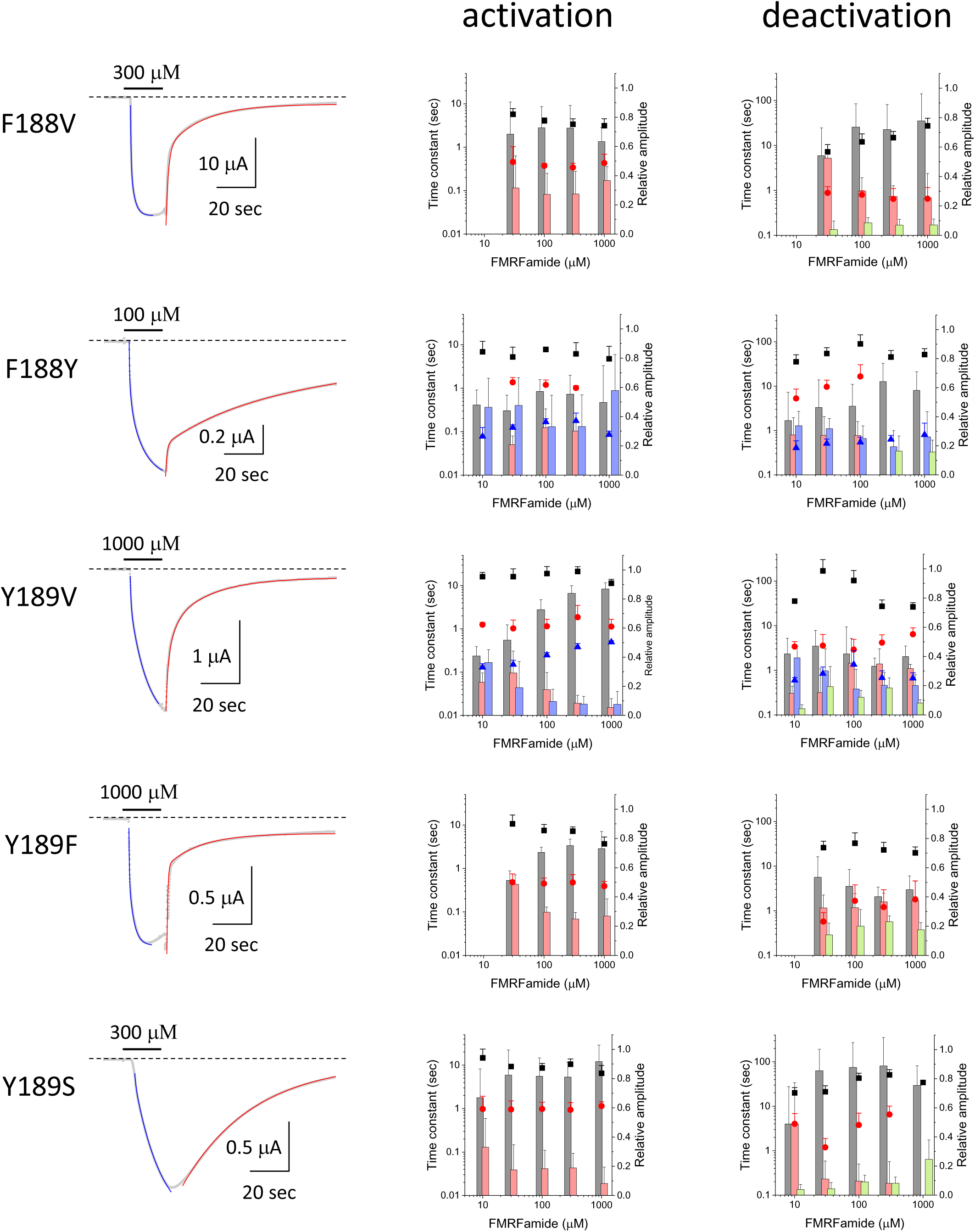
The macroscopic kinetic analysis of F188V, F188Y, Y189V, Y189F and Y189S. The activation was approximated by two (F188V, Y189F, Y189S) or three exponential function (F188Y, Y189V) with a constant and the deactivation was approximated by two (F188V, Y189F, Y189S) or three exponential function (F188Y, Y189V) usually with a constant. The left column shows examples of the fitting. Superimposed blue and red lines are fitted exponential functions. The meanings of symbols and bars for the activation (middle column) and for the deactivation (right column) are as described in the legend of Fig. 6. Number of oocytes was as follows: F188V (5), F188Y (5), Y189V (4–5), Y189F (5), Y189S (8).

**Table 2.**
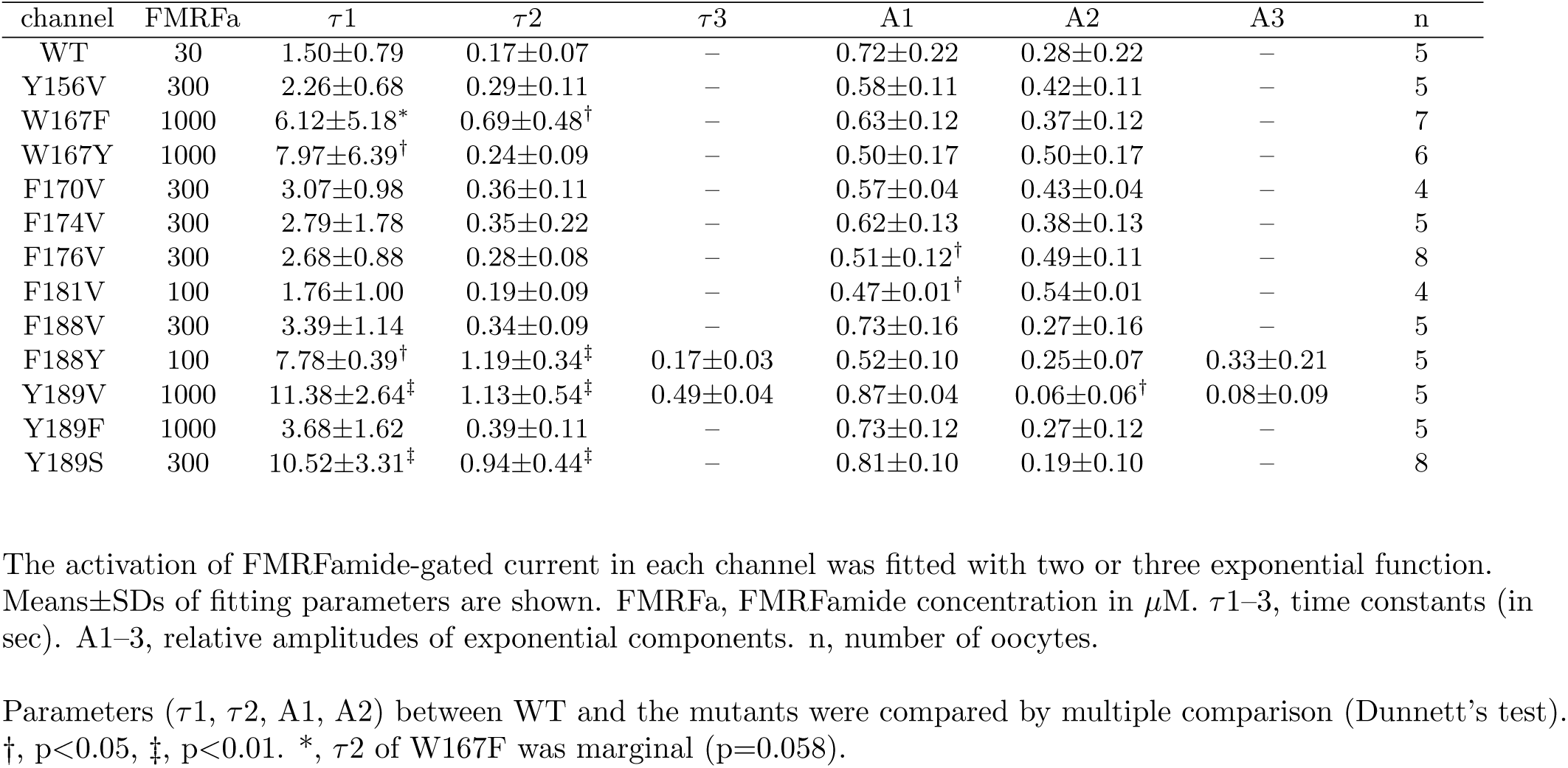
Macroscopic kinetic parameters of the activation of FMRFamide-gated currents

| channel | FMRFa | $\tau_1$ | $\tau_2$ | $\tau_3$ | A1 | A2 | A3 | n |
| --- | --- | --- | --- | --- | --- | --- | --- | --- |
| WT | 30 | 1.50±0.79 | 0.17±0.07 | – | 0.72±0.22 | 0.28±0.22 | – | 5 |
| Y156V | 300 | 2.26±0.68 | 0.29±0.11 | – | 0.58±0.11 | 0.42±0.11 | – | 5 |
| W167F | 1000 | 6.12±5.18* | 0.69±0.48<sup>†</sup> | – | 0.63±0.12 | 0.37±0.12 | – | 7 |
| W167Y | 1000 | 7.97±6.39<sup>†</sup> | 0.24±0.09 | – | 0.50±0.17 | 0.50±0.17 | – | 6 |
| F170V | 300 | 3.07±0.98 | 0.36±0.11 | – | 0.57±0.04 | 0.43±0.04 | – | 4 |
| F174V | 300 | 2.79±1.78 | 0.35±0.22 | – | 0.62±0.13 | 0.38±0.13 | – | 5 |
| F176V | 300 | 2.68±0.88 | 0.28±0.08 | – | 0.51±0.12<sup>†</sup> | 0.49±0.11 | – | 8 |
| F181V | 100 | 1.76±1.00 | 0.19±0.09 | – | 0.47±0.01<sup>†</sup> | 0.54±0.01 | – | 4 |
| F188V | 300 | 3.39±1.14 | 0.34±0.09 | – | 0.73±0.16 | 0.27±0.16 | – | 5 |
| F188Y | 100 | 7.78±0.39<sup>†</sup> | 1.19±0.34<sup>‡</sup> | 0.17±0.03 | 0.52±0.10 | 0.25±0.07 | 0.33±0.21 | 5 |
| Y189V | 1000 | 11.38±2.64<sup>‡</sup> | 1.13±0.54<sup>‡</sup> | 0.49±0.04 | 0.87±0.04 | 0.06±0.06<sup>†</sup> | 0.08±0.09 | 5 |
| Y189F | 1000 | 3.68±1.62 | 0.39±0.11 | – | 0.73±0.12 | 0.27±0.12 | – | 5 |
| Y189S | 300 | 10.52±3.31<sup>‡</sup> | 0.94±0.44<sup>‡</sup> | – | 0.81±0.10 | 0.19±0.10 | – | 8 |

The activation of FMRFamide-gated current in each channel was fitted with two or three exponential function. Means±SDs of fitting parameters are shown. FMRFa, FMRFamide concentration in $\mu\text{M}$ . $\tau_1$ –3, time constants (in sec). A1–3, relative amplitudes of exponential components. n, number of oocytes.
Parameters ( $\tau_1$ , $\tau_2$ , A1, A2) between WT and the mutants were compared by multiple comparison (Dunnett's test). <sup>†</sup>, $p < 0.05$ , <sup>‡</sup>, $p < 0.01$ . \*, $\tau_2$ of W167F was marginal ( $p = 0.058$ ).

**Table 3.**
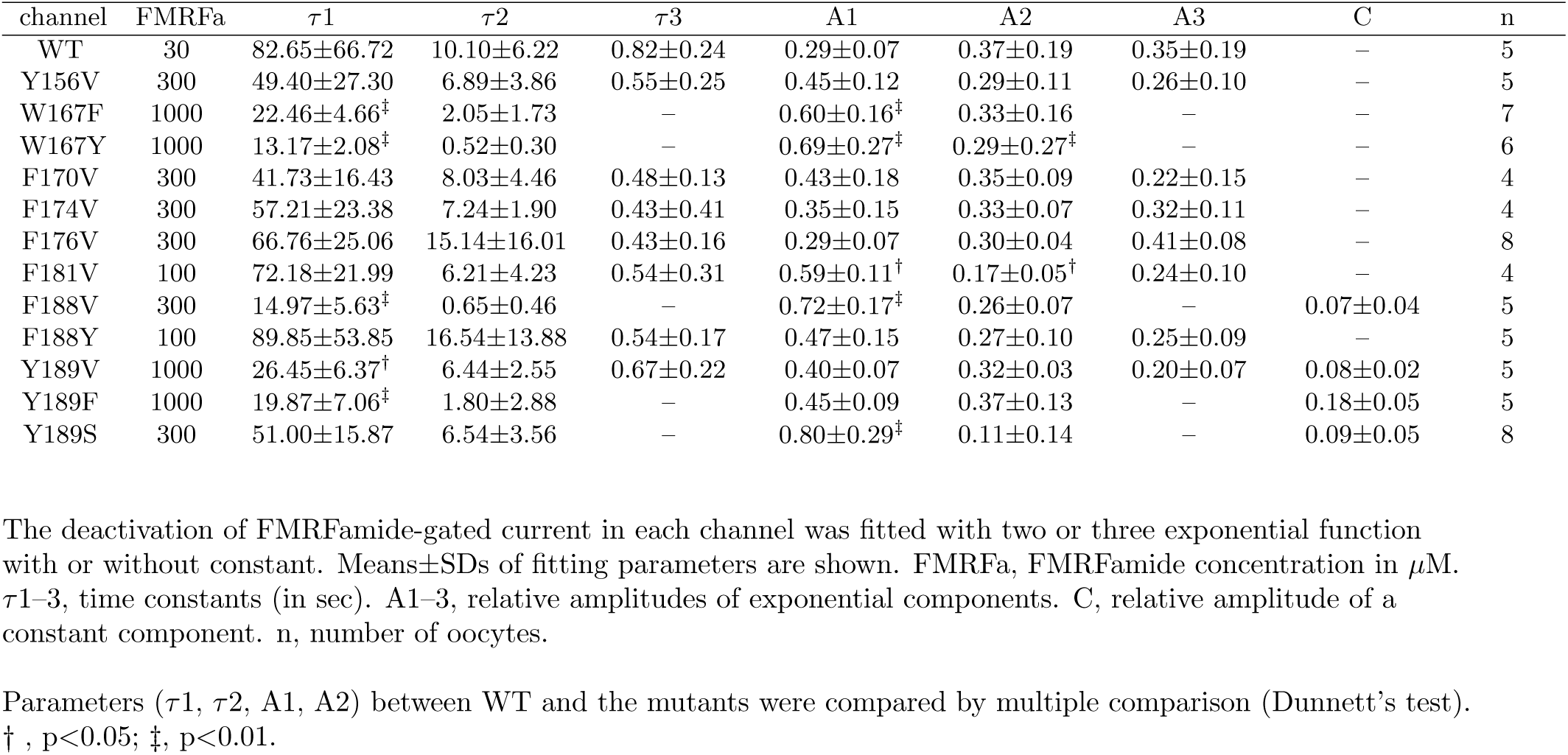
Macroscopic kinetic parameters of the deactivation of FMRFamide-gated currents

| channel | FMRFa | $\tau_1$ | $\tau_2$ | $\tau_3$ | A1 | A2 | A3 | C | n |
| --- | --- | --- | --- | --- | --- | --- | --- | --- | --- |
| WT | 30 | 82.65±66.72 | 10.10±6.22 | 0.82±0.24 | 0.29±0.07 | 0.37±0.19 | 0.35±0.19 | – | 5 |
| Y156V | 300 | 49.40±27.30 | 6.89±3.86 | 0.55±0.25 | 0.45±0.12 | 0.29±0.11 | 0.26±0.10 | – | 5 |
| W167F | 1000 | 22.46±4.66<sup>‡</sup> | 2.05±1.73 | – | 0.60±0.16<sup>‡</sup> | 0.33±0.16 | – | – | 7 |
| W167Y | 1000 | 13.17±2.08<sup>‡</sup> | 0.52±0.30 | – | 0.69±0.27<sup>‡</sup> | 0.29±0.27<sup>‡</sup> | – | – | 6 |
| F170V | 300 | 41.73±16.43 | 8.03±4.46 | 0.48±0.13 | 0.43±0.18 | 0.35±0.09 | 0.22±0.15 | – | 4 |
| F174V | 300 | 57.21±23.38 | 7.24±1.90 | 0.43±0.41 | 0.35±0.15 | 0.33±0.07 | 0.32±0.11 | – | 4 |
| F176V | 300 | 66.76±25.06 | 15.14±16.01 | 0.43±0.16 | 0.29±0.07 | 0.30±0.04 | 0.41±0.08 | – | 8 |
| F181V | 100 | 72.18±21.99 | 6.21±4.23 | 0.54±0.31 | 0.59±0.11<sup>†</sup> | 0.17±0.05<sup>†</sup> | 0.24±0.10 | – | 4 |
| F188V | 300 | 14.97±5.63<sup>‡</sup> | 0.65±0.46 | – | 0.72±0.17<sup>‡</sup> | 0.26±0.07 | – | 0.07±0.04 | 5 |
| F188Y | 100 | 89.85±53.85 | 16.54±13.88 | 0.54±0.17 | 0.47±0.15 | 0.27±0.10 | 0.25±0.09 | – | 5 |
| Y189V | 1000 | 26.45±6.37<sup>†</sup> | 6.44±2.55 | 0.67±0.22 | 0.40±0.07 | 0.32±0.03 | 0.20±0.07 | 0.08±0.02 | 5 |
| Y189F | 1000 | 19.87±7.06<sup>‡</sup> | 1.80±2.88 | – | 0.45±0.09 | 0.37±0.13 | – | 0.18±0.05 | 5 |
| Y189S | 300 | 51.00±15.87 | 6.54±3.56 | – | 0.80±0.29<sup>‡</sup> | 0.11±0.14 | – | 0.09±0.05 | 8 |

The deactivation of FMRFamide-gated current in each channel was fitted with two or three exponential function with or without constant. Means±SDs of fitting parameters are shown. FMRFa, FMRFamide concentration in $\mu\text{M}$ . $\tau_1$ –3, time constants (in sec). A1–3, relative amplitudes of exponential components. C, relative amplitude of a constant component. n, number of oocytes.
Parameters ( $\tau_1$ , $\tau_2$ , A1, A2) between WT and the mutants were compared by multiple comparison (Dunnett's test).
<sup>†</sup>, $p < 0.05$ ; <sup>‡</sup>, $p < 0.01$ .

In most channels examined in the present study, the activation was reasonably well-fitted by two exponential function but three exponential function was required to describe the deactivation. At 10 *µ*M FMRFamide or higher (i.e., fully activated condition), the fast and slow time constants for the activation of WT were *∼*100–300 msec and *∼*1– 3 sec, respectively (Fig. 6, Table 2). The relative amplitude of slow component was usually larger than the fast one. Although the fast and intermediate time constants for the deactivation were also in the order of sub-seconds to seconds, the slow time constant during the deactivation was *∼*30 sec or more, resulting in the long lasting decaying currents after washing out of the peptide (Fig. 6, Table 3). The slow component tended to be larger at higher concentration of FMRFamide.

The activation kinetics of the most mutants in the *α*1–*α*2 region was more or less similar to that of WT (Figs. 6 and 7, Table 2). A slight but noticeable difference of the macroscopic kinetics was observed in the W167 mutants. In W167F or W167Y, the slow time constant for the activation was *∼*10 sec even at high concentration of FMRFamide, resulting in the slower rising phase of the FMRFamide-gated currents compared to WT and other mutants in the *α*1–*α*2 region (Fig. 6, Table 2). On the other hand, the deactivation kinetics of some mutants (Y156V, F170V, F174V, F181V) seemed to be slightly slower than WT. This is largely due to the increase in amplitude of the slow deactivation component (Figs. 6 and 7, Table 3). At higher concentration of FMRFamide, the slow component of deactivation became more dominant in most mutants (see Figs 6 and 7). The deactivation kinetics of W167F or W167Y was reasonably well approximated by two exponential function with the slow time constant of *∼*10–20 sec which was apparently faster than the slow time constants of WT and other mutants in the *α*1–*α*2 region (Table 3).

By contrast, the F188 and Y189 mutants in the loop between *α*2 and *α*3 changed macroscopic kinetics of FaNaC rather drastically (Fig. 8, Tables 2 and 3). The slow time constants for activation of F188 and Y189 mutants were quite different depending on the mutants but the values were clearly larger than that of WT (Table 2). Although the slow time constants of deactivation in these mutants were similar to or slightly less than that of WT, the relative amplitude of the slow components was larger (Fig. 8, Table 3), resulting in the prolongation of the deactivation phase, especially in F188Y and Y189S. Indeed, the decaying phase of some F188 and Y189 mutants was too slow to fit exponential functions adequately without a constant component.

### Docking simulation in the WT model

The dose-response relationship is a convenient measure of the steady state activation of the ligand-gated ion channels. However, the shift of the dose-response relationship of the ligand-gated ion channel by mutation does not tell us what mechanism is disturbed by the mutation: The shift can be produced by the alteration of the ligand binding reaction and/or the modification of the channel gating [9]. To explore whether the conserved aromatic residues in the finger domain of FaNaC are involved in FMRFamide binding, we carried out the docking simulation using the homology model of AkFaNaC and AutoDock Vina (see Materials and Methods). Of course, the docking site as well as the docking pose of FMRFamide obtained by the simulation may not necessarily be the one which is crucial for the FMRFamide-dependent conformational changes of FaNaC. Nonetheless, the docking simulation may be useful to see whether the docking of FMRFamide is disturbed by a particular mutation.

We restricted the search area of docking simulation to the upper part of the finger domain (see Materials and Methods). A rectangular box shown by black lines in Fig. 9a indicates a search area. The docking poses of FMRFamide were classified by coordinates of the first and fourth phenylalanine of FMRFamide (Phe^1^ and Phe^4^) in the cuboidal space (see Materials and methods). 2000 docking simulations showed rather broad docking profile in the finger domain of FaNaC. First of all, we collected docking poses which appeared more than 20 times during 2000 simulations for further analysis. This criterion discriminated 14 docking types, the total docking counts of which was 817 (*∼*41 % of 2000 simulations). We checked the docking poses by PyMol, lumped similar poses together, and determined five dominant types (*∼*34% of 2000 simulations). All the FMRFamide poses in the five types are superimposed on the WT model in Fig. 9b, showing the docking profile in the simulations. Docking poses of FMRFamide in each type are shown separately in Fig. 9c. The number of superimposed docking poses in each type are as follows: type 1 (169), type 2 (84), type 3 (99), type 4 (99), type 5 (228). Estimated affinities (Δ*G*) by AutoDock vina in the five types were similar and their mean values were as follows (kcal/mol): 1 (-6.3), 2 (-6.5), 3 (-6.4), 4 (-6.5), 5 (-6.5).

The docking poses of FMRFamide in the present simulations were roughly separated into two patterns: one was a folded structure due to aromatic stacking between Phe^1^ and Phe^4^ of FMRFamide (types 3, 4) and the other was an elongated one (types 1, 2, 5). In all the dockings, aromatic-aromatic interactions between FMRFamide and FaNaC were rather clear and specific interactions between Phe^1^ and/or Phe^4^ of FMRFamide and aromatic residues in the finger domain were recognized (Fig. 9c). In the type 1, the aromatic stacking interactions are seen between Phe^1^ and W167 as well as Phe^4^ and F176. In the type 2, the aromatic stacking is observed between Phe^4^ and W167. In the types 3 and 4, FMRFamide is folded by intramolecular aromatic stacking and the benzene rings of Phe^1^ and Phe^4^ are nearly perpendicular to the benzene ring of F181. In the type 5, the aromatic stacking interactions are seen between Phe^1^ and F181 as well as Phe^4^ and F174. Because Y156 is close to F181, Y156 appears to influence the dockings in the types 3–5.

### Comparison of FMRFamide Dockings among the mutant models

We next constructed the mutant models and carried out 2000 docking simulations in each model. We compared the results of docking simulations by counting docking poses which were classified into one of the five types described above (the docking counts are summarized in Supplementary Table 1). If a mutation disturbs the docking of FMRFamide, the count number in any one of the five types is expected to become smaller than that obtained in the WT model. In such cases, other docking poses are increased in the present simulations, which may or may not increase the numbers classified into other types among the five dominant types. The difference between the WT model and the mutant (ΔCount) in the five docking types are illustrated in Fig. 10. ΔCount is a normalized difference of the count numbers of the form, (the docking counts in mutant */* the docking counts in WT) *−* 1. The value should be close to zero if the difference in the number of docking poses classified into one of the five types is small. As shown in Fig. 10, the results in some mutant models were clearly different from the ones obtained in the WT model. In the Y156V model, the docking counts in types 4 and 5 were reduced by 84.9% and 87.3%, respectively. In the W167V model, the docking counts in type 1 was absent and the reduction of the counts was 70.2% in type 2. In the W167F model, the docking counts in types 1 and 2 were reduced by 24.9% and 41.7%, respectively. In the W167Y model, the docking counts in types 1 and 2 were reduced by 98.1% and 27.4%, respectively. The docking counts in type 1 was also absent in the F176V model. In the F174V model, the docking counts in types 3–5 were reduced by 42.4%, 90.9%, 96.9%, respectively. In the F181V model, the docking counts in type 3 was absent and the docking counts in types 4 and 5 were reduced by 96.0% and 50.9%, respectively. Most of these results are explainable by the aromatic interactions between the aromatic residues in FaNaC and FMRFamide (see Discussion). By contrast to the above mentioned mutant models, the F188 and Y189 mutant models as well as the F170V model did not change the docking counts of five dominant types drastically (Fig. 10, Supplementary Table 1).

**Fig. 9.**
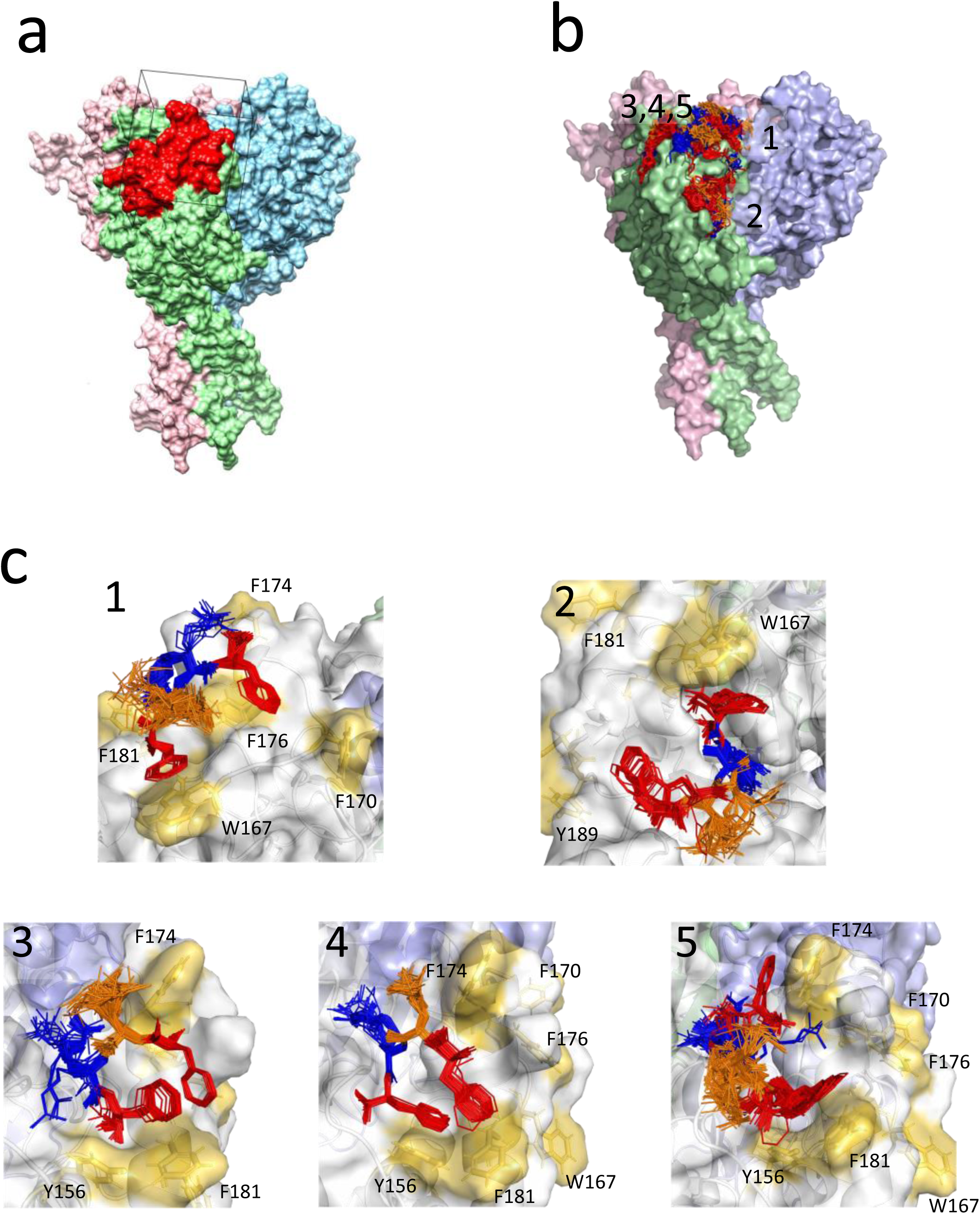
Docking simulations in the WT model. **a:** The WT model showing the search area for the docking simulation by the AutoDock-Vina. The chain A, light green; the chain B, light blue; the chain C, light pink. The finger domain of the chain A is shown by red color. The search area is shown by a black rectangular box. **b:** Docking of FMRFamide on the WT model. All the docking results of the five dominant types extracted from 2000 simulations are superimposed (see Results). The docking positions of the five types are indicated by number. FMRFamide is shown by stick model. Phenylalanine, methionine and arginine of FMRFamide are shown by red, orange and blue colors, respectively. **c:** FMRFamide dockings of the five dominant types. FMRFamide poses classified into one of the five dominant types are superimposed. The chains A, B and C of FaNaC are shown by light gray, light blue and light green, respectively. The conserved aromatic residues are shown by gold.

**Fig. 10.**
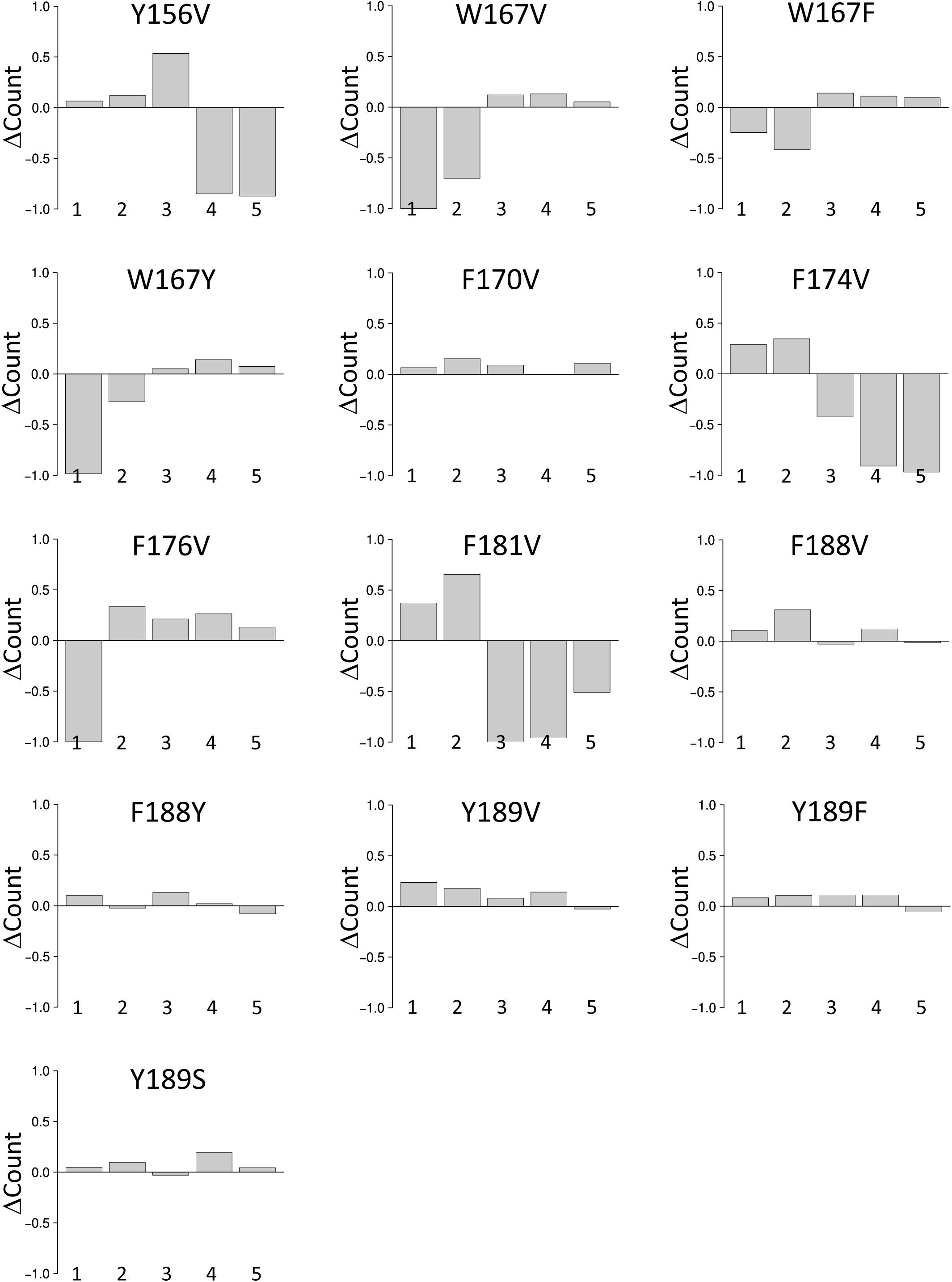
Comparison of the number of docking poses between WT and the mutants. 2000 docking simulations were carried out in the mutant models and the docking poses which were classified into one of the five types described in Fig. 9 were counted. In each graph, ΔCount (see Results) of the mutant in each docking type is illustrated.

## Discussion

Cottrell’s group was the first to demystify the FMRFamide-dependent activation of FaNaC. By using comparative physiological approach, a short stretch of sequence (RRMYFNN in HaFaNaC) was revealed to be important for the FMRFamide recognition of FaNaC [11, 13]. Later, Niu et al [36] have shown that a tyrosine (Y131) in the sequence is a key residue. These results as well as the classical structure-activity relationship of FMRFamide analogue peptides in FaNaC [10] prompted us to consider a hypothesis in which the aromatic-aromatic interaction between FMRFamide and FaNaC is involved in the activation of FaNaC. In the present study, we examined whether the conserved aromatic amino acids in the finger domain of molluscan FaNaCs are involved in the FMRFamide-dependent activation of AkFaNaC by functional analysis of the mutants and computational docking simulations.

The substitution of the conserved aromatic residues in the *α*1–*α*2 region of the finger domain of FaNaC to valine induced *∼*10–50 fold reduction of the potency of FMRFamide to activate the channel (see Table 1). Hill coefficients of some mutants may also appear to change slightly in some mutants although the results of multiple comparison were negative (Table 1). Although the functionality of the aromatic moieties at these positions must be examined more systematically in future, the present study suggests that the conserved aromatic residues in the finger domain of FaNaC are important for the FMRFamide-dependent activation. Among the conserved aromatic residues in the *α*1–*α*2 region, W167 appears to be the most crucial residue for the activation of FaNaC because a single mutation at this site (W167V) totally disrupts the activation of FaNaC by FMRFamide. Because conservative mutants (W167F, W167Y) express the FMRFamide-gated currents, the aromatic moiety at position 167 seems to be critical for the FMRFamide-dependent activation of FaNaC.

In the *α*2–*α*3 loop of the finger domain of molluscan FaNaCs, two tandem aromatic residues, F188 and Y189, are conserved. Substitution of either F188 or Y189 by valine reduced the FMRFamide potency and a double mutation (F188VY189V) made AkFaNaC non-functional, suggesting that tandem aromatic residues in the *α*2–*α*3 loop are critically important for the FMRFamide-dependent activation. Indeed, Y189V showed the largest reduction of the potency of FMRFamide among the functional mutants tested in the present study. As described in Results, the dose-response-relationship of Y189V was actually comprised of two components, one of which showed comparable EC50 to WT, implying that the valine-substitution of Y189 may not necessarily deteriorate the binding affinity of FMRFamide but may reduce the efficacy of gating strongly. Both Y189F and Y189S showed better potencies of FMRFamide compared to Y189V, suggesting that aromatic moiety as well as a hydroxyl group at position 189 are necessary for the efficient activation of FaNaC.

The macroscopic activation/deactivation kinetics of FaNaC was at most modestly affected by the mutations of aromatic residues in the *α*1–*α*2 region. Although the activation was rarely affected by the most mutants in this region, the deactivation was slightly retarded. Such retarded deactivation became more prominent at higher concentration of FMRFamide (Figs. 6 and 7). The slow deactivation was most prominent in F181V among the mutants in this region. The W167 mutants were again unique because they showed slightly slower activation and faster deactivation compared to WT. Some kinetic difference between WT and the mutants in the *α*1–*α*2 region raises a possibility that the kinetic modifications may affect the FMRFamide dose-response relationships in these mutants. More substantial modification of the macroscopic kinetics was observed in F188 and Y189 mutants. The activation rates of F188 and Y189 mutants were in general slower than WT, which was prominent in F188Y, Y189V and Y189S (Fig. 8, Table 2). The deactivation rates of these mutants were also slower because the slow component dominated the deactivation (Fig. 8, Table 3). These results clearly suggest that the mutation at position 188 and 189 affect the gating properties of FaNaC, which could be the main cause for the low potency of FMRFamide in these mutants.

Recent comparative analysis of FaNaC by Dandamudi et al [14] have shown that several molluscan FaNaCs previously not examined are also activated only by FMRFamide and its close analogues (FLRFamide, FMKFamide) but rarely by other invertebrate peptides, extending the previous observations [10]. More surprisingly, they have shown that annelid FaNaCs, *Capitella* FaNaC and *Malacoceros* FaNaC, are activated by FVRIamides which are ineffective in molluscan FaNaCs [14]. Also, the annelid FaNaCs seem to be more sensitive to FMRFamide because their EC50s for FMRFamide are *∼*10 times smaller than the EC50s of molluscan FaNaCs [14]. As described above, W167 and Y189 are important aromatic amino acids for the FMRFamide-dependent activation in AkFaNaC. These two aromatic residues are indeed strictly conserved in the molluscan FaNaCs (Supplementary Fig. 3). In the annelid FaNaCs, however, W167 and Y189 are replaced to tyrosine and serine, respectively (Supplementary Fig. 3). Moreover, the *α*1–*α*3 region of the finger domain is longer in annelid FaNaCs and contains more aromatic residues (Supplementary Fig. 3). It can be imagined that the phylogenetic divergence of the *α*1–*α*3 region, especially the additional aromatic residues, has changed the functionality of FaNaC.

In the present heuristic docking simulations carried out in the WT model, we found some of the conserved aromatic residues can interact with 1st and/or 4th phenylalanine of FMRFamide (Y156, W167, F174, F176, F181, see Fig. 9). Although our docking simulations are certainly limited by the homology model in the present study, the results show that the docking poses prominent in the WT model are less abundant in some mutants of the conserved aromatic residues in the *α*1–*α*2 region, consistent with a notion that the aromatic residues are involved in the binding sites of FMRFamide. Some of the presumed binding sites of FMRFamide revealed by the docking simulations, however, do not appear to be related to the activation of FaNaC. Some docking poses (types 3–5) were absent or greatly decreased in the F181V model in spite of the fact that the FMRFamide potency in F181V was almost the same to that in WT. The inconsistency between functional and computational results retracts a possibility that the docking poses in types 3–5 contribute to the activation of FaNaC directly. Because the docking simulations in the Y156V and F174V models showed similar reduced dockings in types 3–5, the low potency of FMRFamide in Y156V or F174V is not readily explained by the results of docking simulations.

On the other hand, the results in the W167 mutant models are more consistent with functional experiments. As described, W167V is the only point mutant which abolishes the FMRFamide-dependent gating of FaNaC in the present study. In the docking simulations, *π*-*π* stacking between W167 and Phe of FMRFamide is seen in types 1 and 2. These docking poses were much reduced in the W167V model, suggesting that the *π*-*π* stacking are necessary for the dockings. Conservative mutants, W167F and W167Y, were functional but their EC50s were larger than that of WT. In the W167F and W167Y models, the docking counts in types 1 and 2 were also decreased but to a lesser extent. These results are consistent with a notion that the aromatic moiety at position 167 is involved in the FMRFamide binding which is critical for the activation of FaNaC.

By contrast to the docking results of above mentioned models, the docking results in F188 and Y189 mutant models were almost identical to that obtained in the WT model, consistent with a notion that the functional changes observed in F188 and Y189 mutants are due to the deterioration of the gating efficacy of FaNaC rather than the FMRFamide binding. Our results, however, do not exclude a possibility that F188/Y189 is involved in the FMRFamide binding. Because the loop connecting *α*2 and *α*3 in which F188 and Y189 are reside is close to a presumed docking site involving W167 (ex., type 2 in Fig. 9c), some interactions between F188/Y189 and the bound FMRFamide are likely during the activation gating.

In the present homology models, the C-terminal half of specific insertion I of FaNaC which is in the loop structure in the upper thumb domain was removed (see Materials and methods). Altoughh HaFaNaC tolerates the deletion of C-terminal half of the specific insertion I, the full deletion of the specific insertion I makes HaFaNaC non-functional, suggesting that N-terminal half of the specific insertion I in the thumb domain is important for the FMRFamide-dependent activation [36]. Because the upper thumb domain is close to the *α*1–*α*3 region of the finger domain, a proposed FMRFamide binding involving W167 may promote the functional linkage between the *α*1–*α*3 region of the finger domain and the specific insertion I in the thumb domain, which may be necessary for the activation gating of FaNaC. Of course, these possibilities must be rigorously examined in future experiments but some functional coupling between the upper finger domain and the thumb domain of FaNaC has been already suggested [36].

The hypothesis for the activation of FaNaC described above is well in accord with the activation motion suggested in other DEG/ENaC channels. Numerous structural and functional studies in ASIC indicate that the interaction among the finger, thumb and *β*-ball domains is involved in the activation as well as the desensitization of the channels [6, 26, 32, 45, 51, 52]. Psalmotoxin (PcTx1) inhibits the H^+^-activated transient current or evokes the steady current in ASIC [8]. PcTx1 interacts with the finger and thumb domains (acidic pocket) and binds between the interfaces of subunits, which inhibits the ASIC currents [2, 15]. Texas coral snake toxin (MitTx) opens the ASIC pore [5] by binding to the thumb domain [3]. The information about the gating modifier toxins is particularly interesting because the binding regions of these toxins in ASIC are close to type 2 docking site in the WT model described above. The conserved aromatic residues in the finger domain of FaNaC are unique in FaNaC and rare or absent in other DEG/ENaC channels. The aromatic-aromatic interactions between FMRFamide and the finger domain of FaNaC may reduce the energetic barrier for the interactions among the finger, thumb and *β*-ball domains in FaNaC, leading to the activation gating.

ASIC is also known to be modulated by FMRFamide and related peptides (RFamide) [1, 7, 50]. RFamide does not activate ASIC but a fast desensitization of ASIC is retarded by pretreatment with FMRFamide. Recent computational and mutagenic analysis for the binding site(s) of RFamide in ASIC have revealed that the central vestibule of ASIC is a most probable binding region of the peptide [4, 41], which is consistent with a quite slow peptide association rate in ASIC (*∼*1 min pre-application of RFamide is necessary to obtain a maximum modulation in ASIC) [4]. Recently, Dandamudi et al [14] have identified seven residues (F231, I281, R288, G316, R324, S447, R529) in AkFaNaC, mutation of which has made AkFaNaC unresponsive to 100 *µ*M FMRFamide. The identified seven residues may be involved in the FMRFamide binding sites although the other explanation is equally possible as clearly stated in their paper [14]. A drawback of the binding site hypothesis is that the most of seven residues are in the palm domain and not exposed to the outside of extracellular domain of FaNaC. If they are involved in the FMRFamide binding for activation, the peptide binding rate is expected to be slow as shown in the case of ASIC, which is not consistent with the activation rate of FMRFamide-gated currents (see Fig. 6, Table 2) [24].

In summary, we have examined the conserved aromatic residues in the finger domain of FaNaC and found that some of them are critically involved in the FMRFamide binding and/or the activation gating. Although the FMRFamide binding sites are still obscure, the present results are in accord with the previous studies in FaNaC [11, 13, 36] and reinforce a hypothesis that FMRFamide binding sites are in the finger domain. Certainly much more work needs to be done to better understand the functionality of the identified aromatic residues and clarify the binding-gating problem of the peptide-gated Na^+^ channel.

## Acknowledgment

This work was supported by JSPS KAKENHI Grant Number JP15K07149.

## Disclosures

No conflicts of interest, financial or otherwise, are declared by the authors.

## Authors’ contributions

YF conceived research; YF designed experiments; YF and IT made mutant channels; IT performed electophysiological experiments by TEVC. YF performed electrophysiological experiments by COVC; IT and YF analyzed data; YF performed the homology modeling and the docking simulations. YF wrote a paper; IT and YF approved the final version of the paper.

## Notes

### Competing Interest Statement

The authors have declared no competing interest.

## References

[1] Askwith CC, Cheng C, Ikuma M, Benson C, Price MP, Welsh MJ (2000) Neuropeptide FF and FMRFamide potentiate acid-evoked currents from sensory neurons and proton-gated DEG/ENaC channels. Neuron, 26:133–141, DOI: 10.1016/s0896-6273(00)81144-7

[2] Baconguis I, Gouaux E (2012) Structural plasticity and dynamic selectivity of acid-sensing ion channel-spider toxin complexes. Nature, 489(7416):400–405, DOI: 10.1038/nature11375

[3] Baconguis I, Bohlen CJ, Goehring A, Julius D, Gouaux E (2014) X-ray structure of acid-sensing ion channel 1-snake toxin complex reveals open state of a Na^+^-selective channel. Cell, 156(4):717–729, DOI: 10.1016/j.cell.2014.01.011

[4] Bargeton B, Iwaszkiewicz J, Bonifacio G, Roy S, Zoete V, Kellenberger S (2019) Mutations in the palm domain disrupt modulation of acid-sensing ion channel 1a currents by neuropeptides. Sci Rep, 9(1):2599, DOI: 10.1038/s41598-018-37426-5

[5] Bohlen CJ, Chesler AT, Sharif-Naeini R, Medzihradszky KF, Zhou S, King D, Sánchez EE, Burlingame AL, Basbaum AI, Julius D (2011) A heteromeric Texas coral snake toxin targets acid-sensing ion channels to produce pain. Nature, 479 (7373):410–414, DOI: 10.1038/nature10607

[6] Bonifacio G, Lelli CI, Kellenberger S (2014) Protonation controls ASIC1a activity via coordinated movements in multiple domains. J. Gen. Physiol., 143(1):105–118, DOI: 10.1085/jgp.201311053

[7] Catarsi S, Babinski K, Seguela P (2001) Selective modulation of heteromeric ASIC proton-gated channels by neuropeptide FF. Neuropharmacology, 41(5):592–600, DOI: 10.1016/s0028-3908(01)00107-1

[8] Chen X, Kalbacher H, Gründer S (2006) Interaction of acid-sensing ion channel (ASIC) 1 with the tarantula toxin psalmotoxin 1 is state dependent. J. Gen. Physiol., 127(3):267–276, DOI: 10.1085/jgp.200509409

[9] Colquhoun D (1998) Binding, gating, affinity and efficacy: the interpretation of structure-activity relationships for agonists and of the effects of mutating receptors. Br. J. Pharmacol., 125(5):924–947, DOI: 10.1038/sj.bjp.0702164

[10] Cottrell GA (1997) The first peptide-gated ion channel. J. Exp. Biol., 200(Pt 18): 2377–2386, DOI: 10.1242/jeb.200.18.2377

[11] Cottrell GA (2005) Domain near TM1 influences agonist and antagonist responses of peptide-gated Na^+^ channels. Pflügers Arch., 450(3):168–177, DOI: 10.1007/s00424-005-1385-7

[12] Cottrell GA, Green KA, Davies NW (1990) The neuropeptide Phe-Met-Arg-Phe-NH_2_ (FMRFamide) can activate a ligand-gated ion channel in *Helix* neurones. Pflügers Arch., 416(5):612–614, DOI: 10.1007/BF00382698

[13] Cottrell GA, Jeziorski MC, Green KA (2001) Location of a ligand recognition site of FMRFamide-gated Na^+^ channels. FEBS Lett., 489(1):71–74, DOI: 10.1016/s0014-5793(01)02081-6

[14] Dandamudi M, Hausen H, Lynagh T (2022) Comparative analysis defines a broader FMRFamide-gated sodium channel family and determinants of neuropeptide sensitivity. J Biol Chem, 298(7):102086, DOI: 10.1016/j.jbc.2022.102086

[15] Dawson RJ, Benz J, Stohler P, Tetaz T, Joseph C, Huber S, Schmid G, Hügin D, Pflimlin P, Trube G, Rudolph MG, Hennig M, Ruf A (2012) Structure of the acid-sensing ion channel 1 in complex with the gating modifier Psalmotoxin 1. Nat Commun, 3:936, DOI: 10.1038/ncomms1917

[16] Dürrnagel S, Kuhn A, Tsiairis CD, Williamson M, Kalbacher H, Grimmelikhuijzen CJ, Holstein TW, Gründer S (2010) Three homologous subunits form a high affinity peptide-gated ion channel in *Hydra*. J. Biol. Chem., 285(16):11958–11965, DOI: 10.1074/jbc.M109.059998

[17] Dürrnagel S, Falkenburger BH, Gründer S (2012) High Ca^2+^ permeability of a peptide-gated DEG/ENaC from *Hydra*. J. Gen. Physiol., 140(4):391–402, DOI: 10.1085/jgp.201210798

[18] Feyfant E, Săli A, Fiser A (2007) Modeling mutations in protein structures. Protein Sci., 16(9):2030–2041, DOI: 10.1110/ps.072855507

[19] Fujimoto A, Kodani Y, Furukawa Y (2017) Modulation of the FMRFamide-gated Na^+^ channel by external Ca^2+^. Pflügers Arch., 469(10):1335–1347, DOI: 10.1007/s00424-017-2021-z

[20] Furukawa Y, Miyawaki Y, Abe G (2006) Molecular cloning and functional characterization of the *Aplysia* FMRFamide-gated Na^+^ channel. Pflügers Arch., 451(5): 646–656, DOI: 10.1007/s00424-005-1498-z

[21] Golubovic A, Kuhn A, Williamson M, Kalbacher H, Holstein TW, Grimmelikhuijzen CJ, Gründer S (2007) A peptide-gated ion channel from the freshwater polyp Hydra. J. Biol. Chem., 282(48):35098–35103, DOI: 10.1074/jbc.M706849200

[22] Green KA, Cottrell GA (1999) Block of the *Helix* FMRFamide-gated Na^+^ channel by FMRFamide and its analogues. Journal of Physiology, 519:47–56, DOI: 10.1111/j.1469-7793.1999.0047o.x

[23] Green KA, Cottrell GA (2002) Activity modes and modulation of the peptide-gated Na^+^ channel of *Helix* neurones. Pflügers Arch., 443(5-6):813–821, DOI: 10.1007/s00424-001-0750-4

[24] Gründer S, Ramirez AO, Jekely G (2022) Neuropeptides and degenerin/epithelial Na^+^ channels: a relationship from mammals to cnidarians. J Physiol, online, DOI: 10.1113/JP282309

[25] Hanukoglu I, Hanukoglu A (2016) Epithelial sodium channel (ENaC) family: Phylogeny, structure-function, tissue distribution, and associated inherited diseases. Gene, 579(2):95–132, DOI: 10.1016/j.gene.2015.12.061

[26] Jasti J, Furukawa H, Gonzales EB, Gouaux E (2007) Structure of acid-sensing ion channel 1 at 1.9 Å resolution and low pH. Nature, 449(7160):316–323, DOI: 10.1038/nature06163

[27] Jeziorski MC, Green KA, Sommerville J, Cottrell GA (2000) Cloning and expression of a FMRFamide-gated Na^+^ channel from *Helisoma trivolvis* and comparison with the native neuronal channel. J. Physiol. (Lond.), 526 Pt 1:13–25, DOI: 10.1111/j.1469-7793.2000.00013.x

[28] KellenbergerS, Schild L (2002) Epithelial sodium channel/degenerin family of ion channels: A variety of functions for a shared structure. Physiological Review, 82: 735–767, DOI: 10.1152/physrev.00007.2002

[29] Kleyman TR, Carattino MD, Hughey RP (2009) ENaC at the cutting edge: regulation of epithelial sodium channels by proteases. J. Biol. Chem., 284(31):20447–20451, DOI: 10.1074/jbc.R800083200

[30] Kodani Y, Furukawa Y (2010) Position 552 in a FMRFamide-gated Na^+^ channel affects the gating properties and the potency of FMRFamide. Zool. Sci., 27(5):440– 448, DOI: 10.2108/zsj.27.440

[31] Kodani Y, Furukawa Y (2014) Electrostatic charge at position 552 affects the activation and permeation of FMRFamide-gated Na^+^ channels. J Physiol Sci, 64(2): 141–150, DOI: 10.1007/s12576-013-0303-6

[32] Krauson AJ, Carattino MD (2016) The thumb domain mediates acid-sensing ion channel desensitization. J. Biol. Chem., 291(21):11407–11419, DOI: 10.1074/jbc.M115.702316

[33] Lamiable A, Thévenet P, Rey J, Vavrusa M, Derreumaux P, Tufféry P (2016) PEP-FOLD3: faster *de novo* structure prediction for linear peptides in solution and in complex. Nucleic Acids Res., 44(W1):W449–454, DOI: 10.1093/nar/gkw329

[34] Lingueglia E, Champigny G, Lazdunski M, Barbry P (1995) Cloning of the amiloride-sensitive FMRFamide peptide-gated sodium channel. Nature, 378(6558):730–733, DOI: 10.1038/378730a0

[35] Lingueglia E, Deval E, Lazdunski M (2006) FMRFamide-gated sodium channel and ASIC channels: a new class of ionotropic receptors for FMRFamide and related peptides. Peptides, 27(5):1138–1152, DOI: 10.1016/j.peptides.2005.06.037

[36] Niu YY, Yang Y, Liu Y, Huang LD, Yang XN, Fan YZ, Cheng XY, Cao P, Hu YM, Li L, Lu XY, Tian Y, Yu Y (2016) Exploration of the peptide recognition of an amiloride-sensitive FMRFamide peptide-gated sodium channel. J. Biol. Chem., 291 (14):7571–7582, DOI: 10.1074/jbc.M115.710251

[37] Noreng S, Bharadwaj A, Posert R, Yoshioka C, Baconguis I (2018) Structure of the human epithelial sodium channel by cryo-electron microscopy. Elife, 7:e39340, DOI: 10.7554/eLife.39340

[38] Perry SJ, Straub VA, Schofield MG, Burke JF, Benjamin PR (2001) Neuronal expression of an FMRFamide-gated Na^+^ channel and its modulation by acid pH. J. Neurosci., 21(15):5559–5567, DOI: 10.1523/JNEUROSCI.21-15-05559.2001

[39] Pettersen EF, Goddard TD, Huang CC, Couch GS, Greenblatt DM, Meng EC, Ferrin TE (2004) UCSF Chimera–a visualization system for exploratory research and analysis. J Comput Chem, 25(13):1605–1612, DOI: 10.1002/jcc.20084

[40] R Development Core Team. *R: A Language and Environment for Statistical Computing*. R Foundation for Statistical Computing, Vienna, Austria, 2011. URL http://www.R-project.org/. ISBN 3-900051-07-0.

[41] Reiners M, Margreiter MA, Oslender-Bujotzek A, Rossetti G, Gründer S, Schmidt A(2018) The Conorfamide RPRFa Stabilizes the Open Conformation of Acid-Sensing Ion Channel 3 via the Nonproton Ligand-Sensing Domain. Mol Pharmacol, 94(4): 1114–1124, DOI: 10.1124/mol.118.112375

[42] S^̌^ali A, Blundell TL (1993) Comparative protein modelling by satisfaction of spatial restraints. J. Mol. Biol., 234(3):779–815, DOI: 10.1006/jmbi.1993.1626

[43] Schmidt A, Bauknecht P, Williams EA, Augustinowski K, Gründer S, Jékely G (2018) Dual signaling of Wamide myoinhibitory peptides through a peptide-gated channel and a GPCR in *Platynereis*. FASEB J., 32(10):5338–5349, DOI: 10.1096/fj.201800274R

[44] Schrödinger, LLC. The PyMOL molecular graphics system, version 1.8. November 2015.

[45] Sherwood TW, Frey EN, Askwith CC (2012) Structure and activity of the acid-sensing ion channels. *Am. J. Physiol.*, Cell Physiol., 303(7):699–710, DOI: 10.1152/ajpcell.00188.2012

[46] Stefani E, Bezanilla F (1998) Cut-open oocyte voltage-clamp technique. Methods Enzymol, 293:300–318, DOI: 10.1016/s0076-6879(98)93020-8

[47] Taglialatela M, Toro L, Stefani E (1992) Novel voltage clamp to record small, fast currents from ion channels expressed in *Xenopus* oocytes. Biophys J, 61(1):78–82, DOI: 10.1016/S0006-3495(92)81817-9

[48] Trott O, Olson AJ (2010) AutoDock Vina: improving the speed and accuracy of docking with a new scoring function, efficient optimization, and multithreading. J Comput Chem, 31(2):455–461, DOI: 10.1002/jcc.21334

[49] Velankar S, Alhroub Y, Alili A, Best C, Boutselakis HC, Caboche S, Conroy MJ, Dana JM, van Ginkel G, Golovin A, Gore SP, Gutmanas A, Haslam P, Hirshberg M, John M, Lagerstedt I, Mir S, Newman LE, Oldfield TJ, Penkett CJ, Pineda-Castillo J, Rinaldi L, Sahni G, Sawka G, Sen S, Slowley R, Sousa da Silva AW, Suarez- Uruena A, Swaminathan GJ, Symmons MF, Vranken WF, Wainwright M, Kleywegt GJ (2011) PDBe: Protein Data Bank in Europe. Nucleic Acids Res., 39(Database issue):D402–410, DOI: 10.1093/nar/gkq985

[50] Xie J, Price MP, Wemmie JA, Askwith CC, Welsh MJ (2003) ASIC3 and ASIC1 mediate FMRFamide-related peptide enhancement of H^+^-gated currents in cultured dorsal root ganglion neurons. Journal of Neurophysiology, 89:2459–2465, DOI: 10.1152/jn.00707.2002

[51] Yang H, Yu Y, Li WG, Yu F, Cao H, Xu TL, Jiang H (2009) Inherent dynamics ofthe acid-sensing ion channel 1 correlates with the gating mechanism. PLoS Biol., 7 (7):e1000151, DOI: 10.1371/journal.pbio.1000151

[52] Yoder N, Yoshioka C, Gouaux E (2018) Gating mechanisms of acid-sensing ion channels. Nature, 555(7696):397–401, DOI: 10.1038/nature25782

[53] Zhainazarov AB, Cottrell GA (1998) Single-channel currents of a peptide-gated sodium channel expressed in *Xenopus* oocytes. J. Physiol. (Lond.), 513 (Pt 1): 19–31, DOI: 10.1111/j.1469-7793.1998.019by.x Legends of Figures

