## Supplemental information for "Aromatic amino acids in the finger domain of the FMRFamide-gated Na^+^ channel are involved in the FMRFamide recognition and the activation"

Supplementary figures and tables

Table 1
Figure 1
Figure 2
Figure 3

Table 1. Comparison of the number of docking counts between WT and the mutants

| type | WT | Y156V | W167V | W167F | W167Y | F170V | F174V | F176V | F181V | F188V | F188Y | Y189V | Y189F | Y189S | VV* |
|---|---|---|---|---|---|---|---|---|---|---|---|---|---|---|---|
| 1 | 169 | 180 | 0 | 127 | 3 | 180 | 218 | 0 | 232 | 187 | 186 | 209 | 183 | 177 | 188 |
| 2 | 84 | 94 | 25 | 49 | 61 | 97 | 113 | 112 | 139 | 110 | 82 | 99 | 93 | 92 | 92 |
| 3 | 99 | 152 | 111 | 113 | 104 | 108 | 57 | 120 | 0 | 96 | 112 | 107 | 110 | 96 | 104 |
| 4 | 99 | 15 | 112 | 110 | 113 | 99 | 9 | 125 | 4 | 111 | 101 | 113 | 110 | 118 | 96 |
| 5 | 228 | 29 | 240 | 250 | 245 | 253 | 7 | 258 | 112 | 225 | 210 | 222 | 215 | 238 | 227 |

The number of docking poses classified into the five dominat types in 2000 docking simulations are shown.
*, F188VY189V

**Legends for Supplementary Figures**

**Fig. 1** Check of the perfusion speed. The liquid exchange rate was estimated by the change in junction potential at the tip of blunt microelectrode when the external solution was changed. A: A picture showing the domed membrane of oocyte in the top chamber of COVC system. A left-side black bar (arrow) is a tip of trimmed 27 G needle used as an inlet pipe and a microelectrode is positioned near the rim of the domed membrane. The suction pipette for the outlet is out of sight. The change in junction potential at five positions (labeled 1 to 5) was measured. The position 2 is the top of the domed membrane and others were near the rim. B: Examples of the change in junction potential measured at five positions. The control solution was ND96 and the solution was switched to 100 mM KCl for 10 seconds. C: The time constant of the change in junction potential during the onset of perfusion by 100mM KCl. D: The time constant of the change in junction potential during the offset of perfusion by 100mM KCl. Both the onset and the offset were approximated by a single exponential function. The data were obtained from a single setup with multiple measurements (5 times each).

**Fig. 2** Comparison of homology models of AkFaNaC and its deletion mutant. The closed-state structure of cASIC1 (5WKU) was used as a template. 50 homology models were made by Modeller in both AkFaNaC and WT-del (the deletion mutant of AkFaNaC, see the main text). Because the specific insertion II of FaNaC is totally absent in cASIC, the structure is not well determined and seen as an extended loop in AkFaNaC. Note the helical structure in the thumb domain of AkFaNaC is also different from cASIC. By the deletion of the C-terminal half of the specific insertion I, the overall structure of the thumb domain of WT-del becomes more close to cASIC.

**Fig. 3** Alignment of the finger domains of some FaNaCs. Scientific name and NCBI accession number are as follows: *Aplysia kurodai* (AB206707), *Lottia gigantea* (XP_009046425), *Pomacea canaliculata* (XP_025106863), *Biomphalaria glabrata* (XP_013063507), *Octopus bimaculoides* (KQ425921), *Crassostrea gigas* (XP_011440506), *Mizuhopecten yessoensis* (XP_021342351), *Malacoceros fuliginosus* (ON156821), *Capitella teleta* (KB300602). Aromatic amino acids are shown by white letters in black background. The position of Y156, W167, F170, F174, F176, F181, F188, Y189 in *Aplysia kurodai* are shown above the sequence. Note W167 and Y189 are completely conserved in the molluscan FaNaCs but not in the annelid FaNaCs.

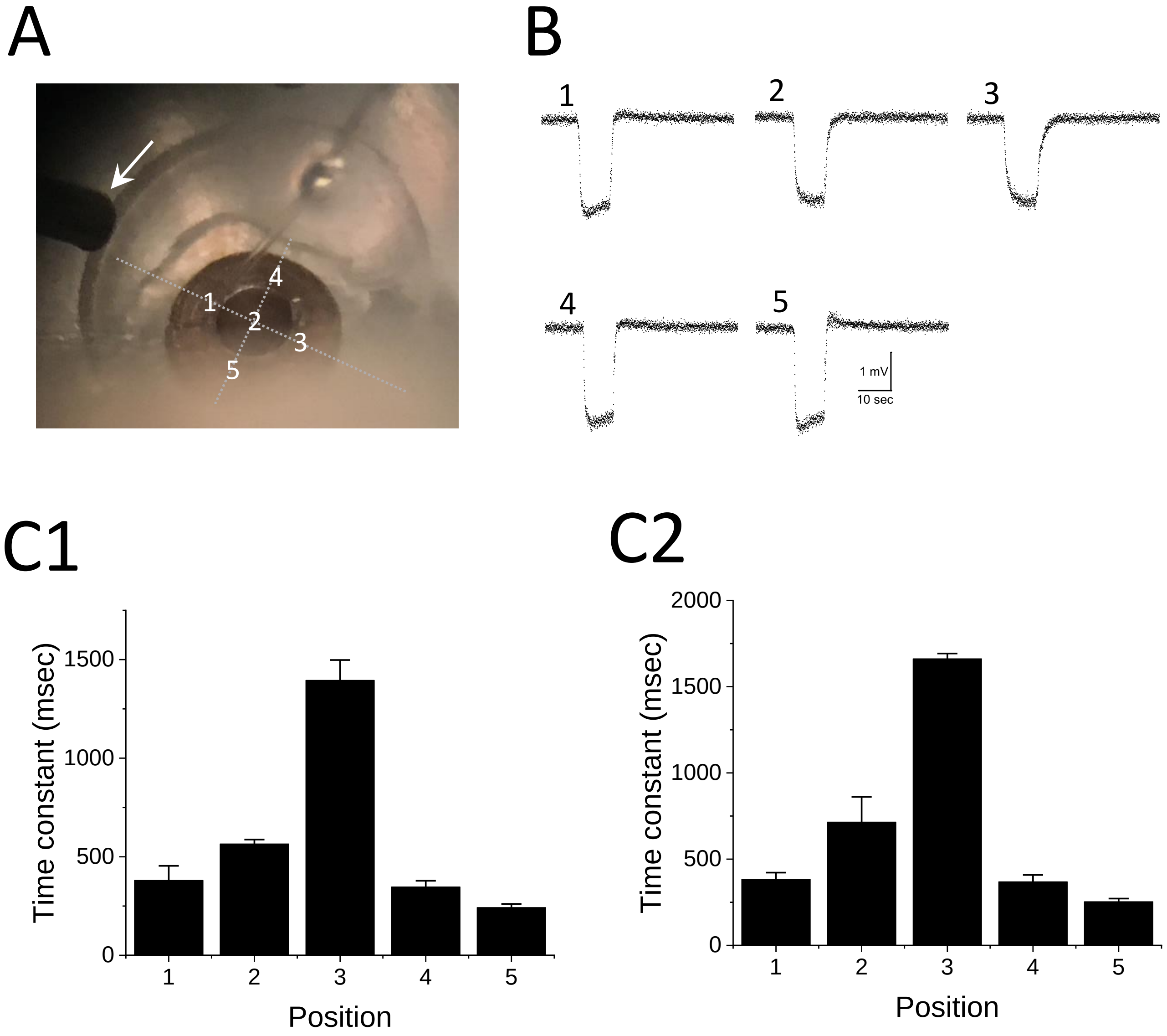
A
1
2
3
4
5
B
1
2
3
4
5
1 mV
10 sec
C1
Time constant (msec)
0
500
1000
1500
1
2
3
4
5
Position
C2
Time constant (msec)
0
500
1000
1500
2000
1
2
3
4
5
Position

Figure 1:

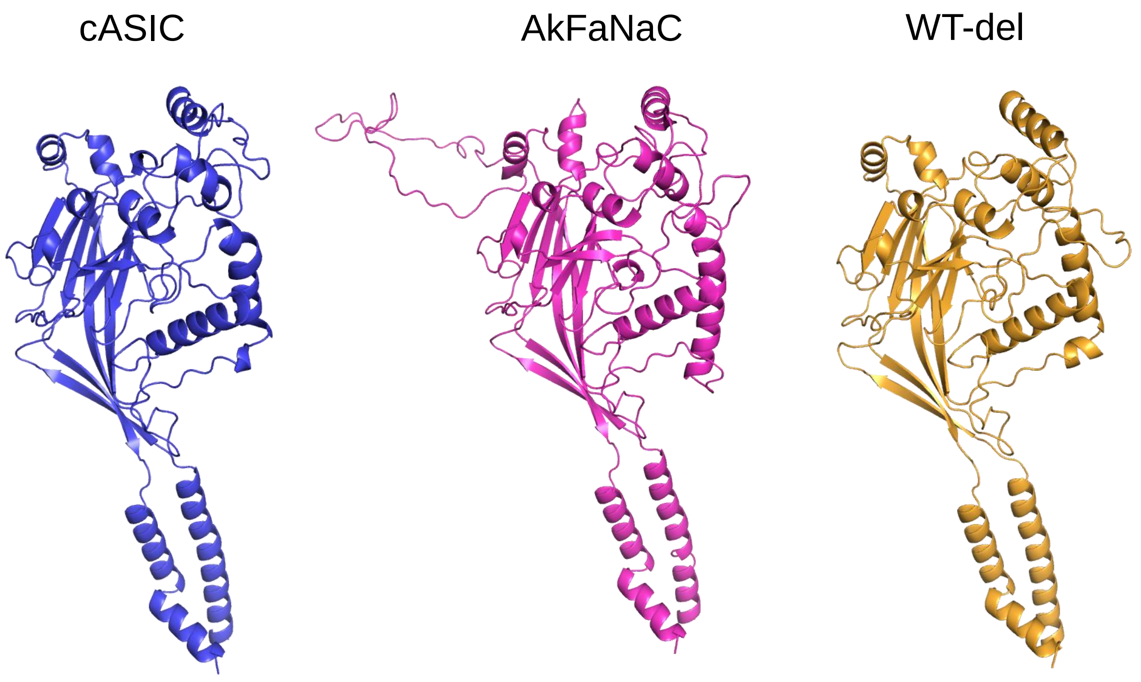


Figure 2:

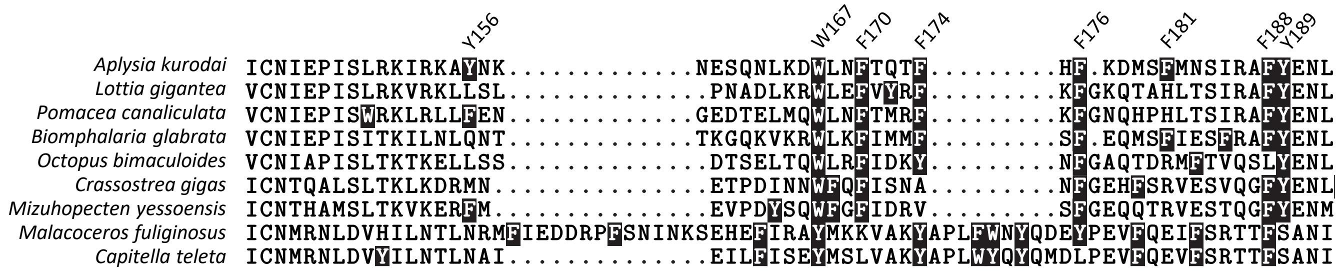


Figure 3:
